# Frizzled-4 regulates β-catenin in endothelial cells exposed to disturbed flow via an atypical Wnt pathway leading to proinflammatory activation and increased permeability

**DOI:** 10.1101/2021.11.12.468370

**Authors:** Matthew Rickman, Mean Ghim, Kuin Tian Pang, Ana Cristina von Huelsen Rocha, Elena M. Drudi, Sureda-Vives, Nicolas Ayoub, Virginia Tajadura-Ortega, Sarah J. George, Peter D. Weinberg, Christina M. Warboys

**Affiliations:** Department of Comparative Biomedical Sciences, Royal Veterinary College, Royal College Street, London, NW1 0TU; Department of Bioengineering, Imperial College London, London, SW7 2AZ; Translational Health Sciences, Bristol Medical School, Research Floor Level 7, Bristol Royal Infirmary, Bristol, BS2 8HW, UK

## Abstract

**Objective:** Endothelial cells are regulated by hemodynamic wall shear stress and multidirectional shear stress is known to promote endothelial dysfunction, although the molecular mechanisms are poorly defined. Wnt pathways play an important role in non-vascular mechanoresponsive cells. Here we investigated their role in endothelial mechanosignalling and endothelial dysfunction.

**Approach & Results:** Human aortic endothelial cells were exposed to shear stress using an orbital shaker. The expression of Frizzled-4 receptors was significantly increased in endothelial cells exposed to low magnitude multidirectional flow (LMMF) relative to high magnitude uniaxial flow (HMUF). Increased expression was also detected in regions of the murine aortic arch exposed to LMMF. The increased Frizzled-4 expression in cultured cells was abrogated following knockdown of R-spondin-3 (RSPO-3) using RNA interference. LMMF also increased the stabilisation and nuclear localisation of β-catenin, an effect that was dependent on Frizzled-4 and RSPO-3. Inhibition of β-catenin using a small molecule inhibitor (iCRT5), or knockdown of Frizzled-4 or R-spondin-3 resulted in a significant reduction of pro-inflammatory gene expression in endothelial cells exposed to LMMF. Stabilisation of the β-catenin destruction complex using IWR-1 under LMMF also reduced pro-inflammatory gene expression, as did inhibition of Wnt5a signalling. Interestingly, inhibition of the canonical Wnt pathway had no effect. Inhibition of β-catenin signalling also reduced endothelial permeability; this was associated with altered junctional and focal adhesion organisation and cytoskeletal remodelling.

**Conclusions:** These data suggest the presence of an atypical Wnt-β-catenin pathway in endothelial cells that promotes inflammatory activation and barrier disruption in response to LMMF.

## Introduction

Endothelial cells (EC) are continuously exposed to hemodymanic wall shear stress (the frictional force per unit area exerted by flowing blood). Surface mechanosensors enable EC to sense and respond to this shear stress via various mechanosignalling pathways^1^. Regulation of these pathways differs greatly depending on local vessel geometry and the resulting shear stress to which EC are exposed, leading in turn to significant differences in EC function. EC in straight, unbranched regions of arteries experience high magnitude, pulsatile but uniaxial wall shear stress that is a key determinant of EC homeostasis, activating cytoprotective signalling pathways and promoting the expression of KLF-2, KLF-4, eNOS and thrombomodulin. Conversely, EC in atherosclerosis-prone areas within regions of high curvature, branching or bifurcation exhibit endothelial dysfunction, with increased permeability, reduced expression of eNOS and activation of pro-inflammatory signalling pathways^2^.

The type of shear stress that leads to the latter behaviour is controversial. Recent work has challenged the consensus that low time average wall shear stress and a high oscillatory shear index drives EC dysfunction and atherogenesis^3^ and instead have highlighted the importance of multidirectional flow (high transverse wall shear stress)^4^; however, the precise trigger is still unknown. The mechanisms by which multidirectional flow influences EC behaviour are poorly defined, although some signalling pathways have been identified that contribute to aspects of flow-dependent EC dysfunction^5–7^. In particular, little attention has been given to the molecular mechanisms and mechanosignalling pathways that promote barrier disruption in EC exposed to multidirectional flow. That is a significant deficiency, given the critical role that elevated permeability plays in the development of cardiovascular disease.

The Wnt signalling pathway is known to regulate responses to mechanical forces in non-vascular mechanoresponsive cells^8^. Transcriptomic analysis reveals that Wnt pathway genes are significantly enriched in EC exposed to atherogenic flow environments^9,10^ suggesting a possible role in promoting flow-mediated EC dysfunction^10^. Wnt signalling pathways (reviewed elsewhere^11^) are highly conserved and require the interaction of Wnt with Frizzled receptors on the cell surface. Complexity arises from the presence of multiple Wnt ligands, Frizzled receptors and co-receptors that can associate in various combinations in a context-dependent manner^12^. Canonical Wnt signalling requires the dephosphorylation and stabilisation of β-catenin, typically through inhibition of the β-catenin destruction complex, allowing β-catenin to translocate to the nucleus and regulate transcription of target genes via interaction with transcription factors e.g. TCF-4^11^. In this study we sought to investigate and dissect the contribution of canonical Wnt signalling pathways in mediating EC dysfunction using an *in vitro* model that provides both high magnitude uniaxial flow (HMUF) and low magnitude multidirectional flow (LMMF)^13^.

## Materials and Methods

### Culture of human aortic endothelial cells and exposure to flow

Human aortic endothelial cells (HAEC) from male and female donors were obtained from Promocell and cultured on fibronectin-coated plasticware in Promocell Endothelial Growth Medium MV with Supplement Mix. Culture medium was replaced every 48-72h and cells were sub-cultured using Trypsin-EDTA solution (0.25%). For flow experiments, cells were seeded (at passage 6-7) in 6-well plates coated with fibronectin (10μg.ml^−1^) and cultured for 24-48h until confluent. For immunostaining experiments, cells were seeded in glass-bottomed 6-well plates (Cellvis). Once confluent, 1.902 ml fresh medium was added to each well (equivalent to 2mm medium height). Plates were placed on an orbital shaker housed inside the incubator (Grant Instruments; 150rpm with 5mm orbital radius) and exposed to flow, induced by the swirling of the media across the base of the well^14–16^. Computational fluid dynamics, using StarCCM+ analysis software, was used to determine various steady-state flow metrics at the base of the well (analysis published elsewhere)^17^. Briefly, EC at the centre of the well were exposed to low magnitude multidirectional flow (LMMF), whereas EC at the edge of the well were exposed to higher magnitude unidirectional flow (HMUF)^13,17^. Cells were exposed to flow for 72h unless stated otherwise.

### Computational fluid dynamics

Flow simulations were carried out with Star CCM+ (version 11.02.009, CD-Adapco, USA). A single well of a 6-well plate was represented as a cylinder of the height of 10 mm and radius of 17.4 mm. The geometry was discretized using a structured cylindrical mesh with 360,000 grid elements. The explicit unsteady model was used. A no-slip condition was imposed at all walls and surface tension was neglected. The top surface of the cylinder was defined as a pressure outlet. The dynamic viscosity and density of medium were 0.78 x 10^3^ Pa.s and 1003 kg/m^3^ respectively. The rotation of the well was modelled by introducing a translating gravitational force with the form

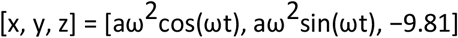

where A is the orbital radius of the shaker, ω is the angular velocity and t is time. The Volume of Fluid model was used to track the free surface of the liquid, which had a height of 2 mm when the well was stationary. Time steps of 1 x 10^−4^ s were each iterated 5 times. Maximum WSS at the base of the well was used to assess convergence. A mesh independence study was performed using 720,000 grid elements, and no difference was observed.

### Transfection with siRNA and TCF reporter plasmid

Prior to seeding into 6-well plates, EC were transfected with siRNA (100 nM) or Cignal™ reporter plasmid (500 ng) by electroporation using a Neon™ Transfection System according to the manufacturer’s instructions. Electroporated cells were seeded directly into wells containing Endothelial Growth Medium (approx. 5 x 10^5^ cells per well) and cultured for 6-8 h until firmly adhered and confluent. Culture medium was replaced before exposure to flow (as above).

### Luciferase Reporter Assay

Following exposure to flow, EC were scraped and aspirated from specific regions of wells to leave either the LMMF or HMUF region only. The remaining cells were lysed with 100 μl Passive Lysis Buffer (Promega) and incubated at room temperature for 15 min with constant agitation. TCF/LEF reporter activity was assessed using a Dual-Luciferase^®^ Reporter Assay System (Promega) according to the manufacturer’s instructions. Luminescence of Firefly (experimental) and *Renilla* (control) luciferase reporters was quantified using a Varioskan Flash Plate Reader. The activity of the Firefly (experimental) reporter was normalised to the activity of the *Renilla* (control) reporter to account for differences in transfection efficiency. Normalised values were also corrected for protein concentration to account for any variations in cell lysis.

### Monocyte adhesion assays

THP-1 monocytes (obtained from ATCC^®^) were cultured in suspension in RPMI-1640 medium supplemented with FBS (10%), L-glutamine (2 mM), Penicillin-Streptomycin (100 IU.ml^−1^) and β-mercaptoethanol (50 μM). Culture medium was replaced every 48-7h and cell density maintained at 8 x 10^5^ - 1 x 10^6^ cells.ml^−1^. Prior to adhesion assays, THP-1 monocytes were incubated with Calcein-AM (1 μg.ml^−1^) for 20 min at 37°C. Following labelling, monocytes were washed, re-suspended in EC growth medium and added to endothelial monolayers for 72h (1 x 10^6^ per well) and incubated at 37°C under static conditions for 60 min. Unbound monocytes were removed by extensive washing with warm EC growth medium and adherent cells fixed using paraformaldehyde (4%) for 5 min. Images of adherent monocytes were captured using an inverted Zeiss Axioplan epifluorescence microscope using a 20× objective with 470/40 nm excitation and 525/50 nm emission filters. The total number of adherent monocytes was captured across 5 fields of view from the centre of each well.

### RNA isolation and qRT-PCR

Following exposure to flow, EC were scraped and aspirated from specific regions of wells to leave either the LMMF or HMUF region only. RLT lysis buffer (Qiagen) supplemented with β-mercaptoethanol was added to wells to lyse cells. In order to obtain enough material, cells from LMMF or HMUF regions from 3 wells were pooled together. RNA was isolated using RNeasy Mini Kits with on-column DNase digestion (Qiagen) according to the manufacturer’s instructions. cDNA was prepared using High-Capacity Reverse Transcription Kit (Thermo Fisher) according to the manufacturer’s instructions. Transcript levels were assessed by quantitative real-time PCR using Fast SYBR Green MasterMix (ABI) and gene-specific primers (Sigma Aldrich) as detailed in Supplementary Table 1. Reactions were carried out using an Applied Biosystems StepOnePlus™ Real-Time PCR System as follows: 95°C for 20 sec, followed by 40 cycles of 95°C for 3 sec and 60°C for 30 sec. All reactions were performed in triplicate and relative expression assessed using the ΔΔCt method using GAPDH as a reference gene.

### Protein extraction and western blotting

Following exposure to flow, EC were scraped and aspirated from specific regions of wells to leave either the LMMF or HMUF region only. Cells were lysed with ice-cold RIPA buffer (SDS (0.1%), Triton-X 100 (1%), sodium deoxycholate (0.5%), NaCl (150 mmol/l), Tris (50mmol/l) at pH 7.4) supplemented with protease and phosphatase inhibitor cocktails (Sigma). In order to obtain enough material, cells from LMMF or HMUF regions from 3 wells were pooled together. Lysates were incubated on ice for 45 min with regular vortexing and centrifuged at 12,000 rpm for 10min to separate soluble and insoluble fractions. For analysis of cytosolic and nuclear fractions, cells were lysed using NE-PER™ Nuclear and Cytoplasmic Extraction Reagent (Thermo Fisher) according to the manufacturer’s instructions. Cells were pooled from LMMF or HMUF regions from 6 wells to obtain enough material. Lysates were analysed by SDS-PAGE and immunoblotting. Membranes were incubated with primary antibodies (Supplementary Table 1) overnight at 4°C and visualised with HRP-conjugated anti-IgG secondary antibodies and enhanced chemiluminescence substrates (Bio-Rad). The expression of PDHX or calnexin was used as a loading control.

### Immunofluorescent staining

Following exposure to flow, EC on glass-bottomed 6-well plates were fixed with 4% paraformaldehyde or ice-cold methanol for 20 min then permeabilised with 0.1% TX-100 for 3 min. After blocking with 5% BSA for 1 h, EC were incubated with primary antibodies (Supplementary Table 1) overnight at 4°C followed by incubation for 1h at room temperature with relevant Alexa Flour 488 or 568-conjugated secondary antibodies diluted 1 in 500 in 1% BSA. EC were incubated with Alexa Fluor-488 Phalloidin (Thermo Fisher) for 20 min at room temperature to visualise the actin cytoskeleton. Nuclei were stained by incubation with DRAQ5 (5 μM). Omission of the primary antibody was used to control for non-specific staining. EC were imaged directly in the well using a Leica SP5 or SP8 Laser Scanning Confocal Microscope. A minimum of 4 fields of view were studied for each flow condition in each well using identical laser power and detector gain settings. Mean fluorescence intensities (MFI) were determined using Leica Imaging Software.

### Quantification of EC permeability

EC were seeded in 6-well plates coated with biotinylated gelatin^18^ and cultured for 72h under static conditions followed by exposure to flow for 72h using the orbital shaker. Medium was replaced with reduced serum (2.5% FBS) EC growth medium (Promocell) for the final 24h of flow exposure. FITC-avidin diluted in reduced serum media (0.38 μM) was added to wells which were incubated at 37°C for 3 minutes. The FITC-avidin solution was then removed and wells washed three times with PBS before fixing with paraformaldehyde (4%) for 10 min. EC were counterstained with VE-cadherin to visualize the distribution of tracer in relation to cell borders. Wells were imaged with a Leica SP5 inverted confocal microscope using a x10, 0.40 NA objective. 9 z-stack images were taken from the LMMF region in the centre of the well (3×3 tile scan) and 12 images from the HMUF regions at the edge of the well. For quantification of FITC-avidin accumulation, a max projection image was computed for each stack of images and the tracer fluorescence was distinguished from background noise by intensity and area thresholding. The resulting binarised image was overlaid on the original image and used as a mask to quantify the FITC-avidin accumulation.

FITC-avidin tracer (0.38 μM) was added to EC monolayers seeded onto biotinylated gelatin matrix, and incubated at 37°C for 3 min^19^. Binding of fluorescently-labelled avidin to biotin occurs underneath, in particular the intercellular junctions^20^, and can be imaged. Monolayers were washed with PBS to remove unbound tracer before fixing with 4% paraformaldehyde for 10 min. Fixed monolayers were then incubated with an anti-VE-cadherin antibody to delineate cell junctions. Stacks of images through the monolayer were obtained from LMMF and HMUF regions using confocal microscopy, and tracer accumulation for each image was computed.

### Immunostaining and imaging of mouse aortas

All animal work was carried out in accordance with Directive 2010/63/EU of the European Parliament on the protection of animals used for scientific purposes, under authorisation of the UK Home Office (Project License No. 70-8934). Male C57BL/6J mice (n=3) were purchased from Charles River Laboratories (Harlow, UK) at 8 weeks of age. Mice were fed a standard chow diet (ad libitum) and maintained on a 12-hour day/night cycle. Mice were sacrificed at 10 weeks of age by terminal anaesthesia using an overdose of pentobarbitone (120 mg/kg i.p). The aorta was fixed and washed by transcardial perfusion with ice-cold saline (0.9%) containing 100 U/ml heparin, followed by perfusion with ice-cold paraformaldehyde (4%). Aortas were removed, further post-fixed overnight in paraformaldehyde (4%) and subject to immunofluorescent staining (as above). Aortas were imaged by *en face* confocal microscopy using a Nikon Spinning Disk Confocal Microscope^21^.

### Statistical Analysis

All data are presented as mean ± standard error of the mean (SEM). A minimum of three independent experiments was carried out for each investigation. Cells from at least three independent donors were used for each experiment. Data were analysed using GraphPad Prism v8.0. A Shapiro Wilk normality test was used to determine whether data were normally distributed. For normally distributed data, statistical significance was determined using a student’s two-tailed t-test when comparing two conditions or by one-way analysis of variance (ANOVA) with Bonferroni’s multiple comparison test when comparing more than one condition. Non-parametric analysis was performed on data that did not pass the normality test; a Mann-Whitney test was used to compare two conditions and a Kruskal-Wallis test with uncorrected Dunn’s test was used to compare multiple conditions (*P<0.05, **P<0.01, ***P<0.001).

## Results

### Frizzled-4 expression is increased by LMMF and promotes endothelial inflammatory signalling

In order to assess the potential flow-dependent regulation of Wnt signalling pathways we investigated the effect of different flow conditions on the expression of Frizzled receptors by using the orbital shaker method. Western blot analysis revealed that LMMF significantly increased the expression of Frizzled-4 relative to HMUF after 24h and that this increase was sustained for 72h of flow exposure (Figure 1a). Demonstrating the potential physiological relevance of this finding, we also observed increased Frizzled-4 receptor in the inner curvature of the aortic arch, a region exposed to chronic flow disturbance^22^, compared to the descending aorta (Figure 1b). To determine whether Frizzled-4 mediates endothelial dysfunction under prolonged LMMF we transfected EC with siRNA targeting Frizzled-4 before exposure to flow. Protein expression of Frizzled-4 was reduced by at least 50% after 48h of flow when compared to EC transfected with scrambled RNA. The expression of Frizzled-5, Frizzled-6 and Frizzled-7 transcript was unaffected (Supplementary Figure S2). Knockdown of Frizzled-4 resulted in significantly reduced expression of several flow-dependent pro-inflammatory genes when compared to scrambled transfected controls (E-selectin, MCP-1, VCAM-1 and IL-8; Figure 1c) and a concomitant reduction in VCAM-1 protein (Figure 1d). Knockdown of Frizzled-4 did not alter the expression of the flow-sensitive genes KLF-2 and eNOS (Supplementary Figure S2) suggesting that EC remain capable of responding to flow.

**Figure 1.**
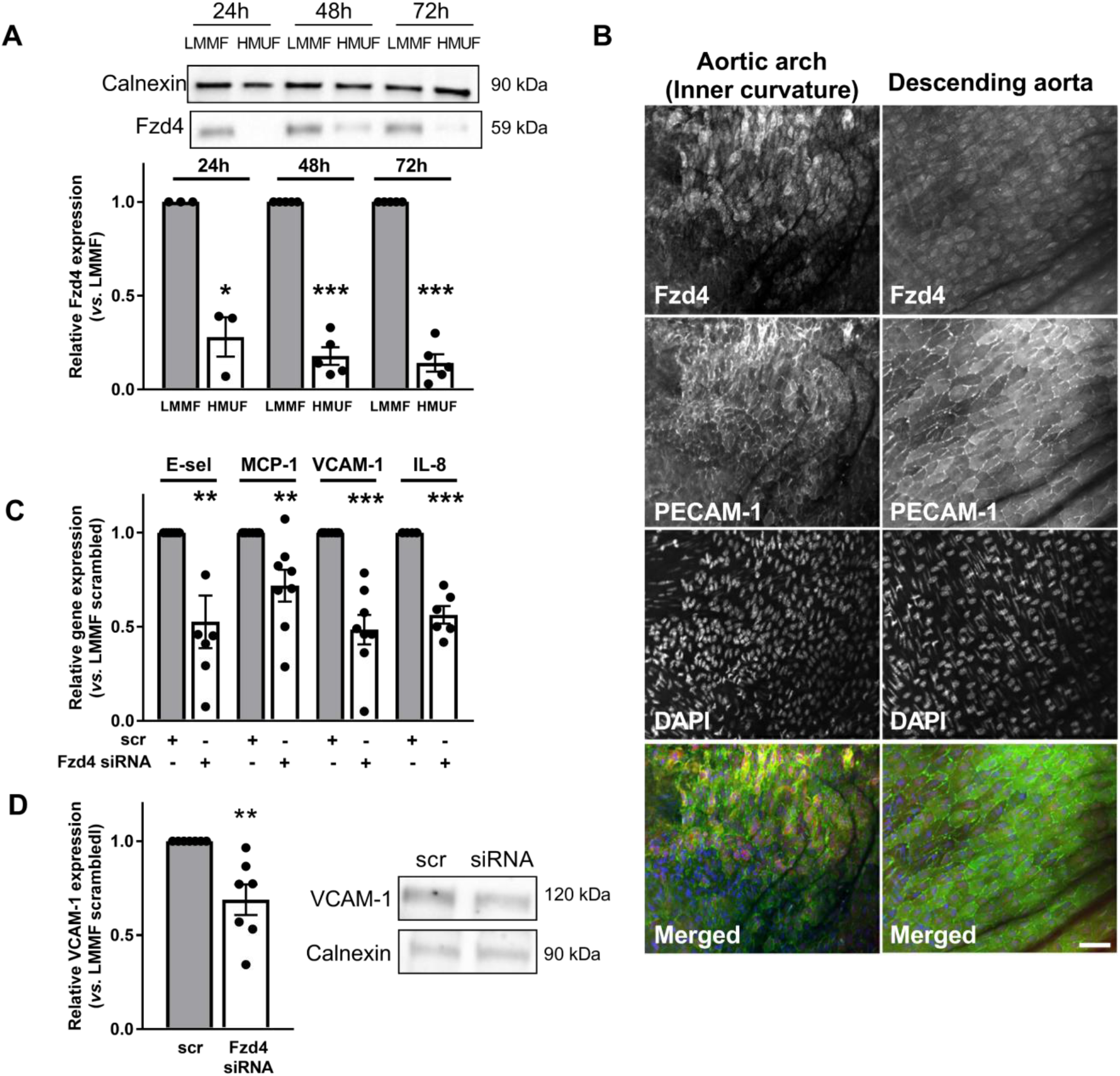
Frizzled-4 is increased in EC exposed to LMMF and regulates pro-inflammatory signalling. **(A)** Protein lysates were obtained from HAEC exposed to LMMF and HMUF for 24-72h. Frizzled-4 expression was analysed by western blot using calnexin as a loading control (n=3-5; analysis by paired t-test at each time point). **(B)** Aortas from C57BL/6 mice were stained with Frizzled-4 and PECAM-1 antibodies and DAPI and imaged *en face* (n=3; representative image shown; scale = 50 μm). **(C)** RNA or **(D)** protein lysates were prepared from EC transfected with Frizzled-4 siRNA or scrambled controls and exposed to LMMF for 48h. **(C)** Gene expression was determined by qRT-PCR using GAPDH as a housekeeping gene (n=6-9; analysis by unpaired t-test). **(D)** Expression of VCAM-1 was assessed by western blot using calnexin as a loading control (n=7; analysis by unpaired t-test; representative blots shown in the panel).

### Frizzled-4 expression under LMMF is regulated by R-spondin-3

We next sought to determine the mechanism by which LMMF elevates Frizzled-4 protein expression. Paradoxically, we found that expression of the Frizzled-4 gene was significantly lower in EC exposed to LMMF relative to HMUF (Figure 2a), pointing towards a post-transcriptional or post-translational mechanism. Since Frizzled-4 protein expression can be regulated by proteasomal degradation, mediated by ZNRF3 ubiquitin ligases, we explored the regulation of this pathway by flow. We found no observable difference between the expression levels of ZNRF3 under LMMF and HMUF (Figure 2b) but we did observe a significant increase in the expression of R-spondin 3 (RSPO-3), which inhibits ZNRF3, in EC exposed to LMMF at both the gene (Figure 2c) and protein level (Figure 2d). The expression of RSPO-1 −2 and −4 was undetectable in these cells (data not shown). These data are consistent with the LMMF-dependent increase in Frizzled-4 being caused by an increased expression of RSPO-3, which would inhibit its proteasomal degradation. Supporting this view, Frizzled-4 gene and protein expression was significantly reduced in EC under LMMF conditions following knockdown of RSPO-3 (Figure 2f). Moreover, knockdown of RSPO-3 also reduced the expression of VCAM-1 in EC exposed to LMMF (Figure 2g), mirroring the effects of Fzd4 knockdown.

**Figure 2.**
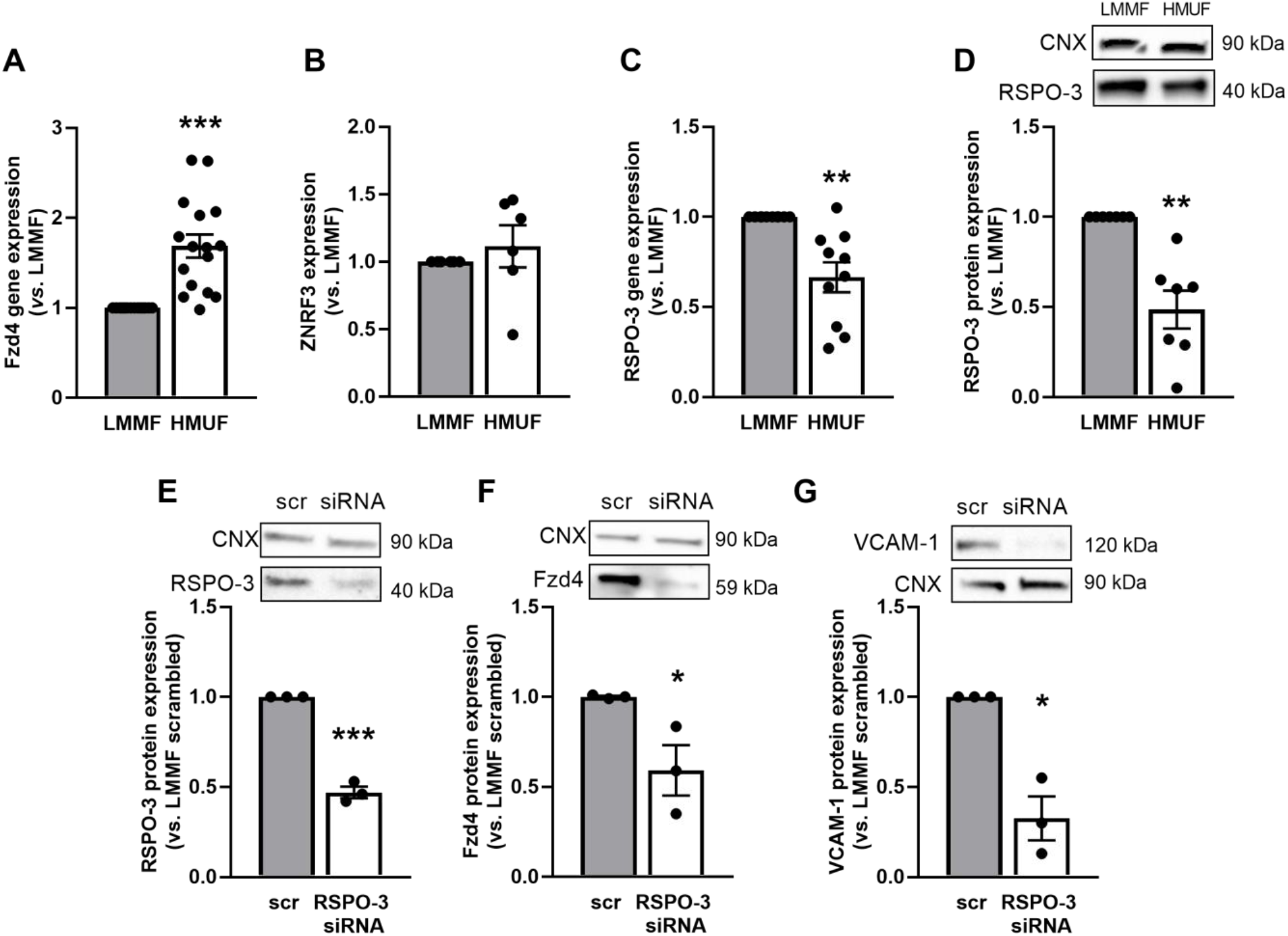
RSPO-3 is increased in EC exposed to LMMF and regulates the expression of Frizzled-4. **(A-D)** Lysates were obtained from HAEC exposed to LMMF and HMUF for 72h. **(A-C)** RNA lysates were prepared and the expression of **(A)** Frizzled-4 (n=16), **(B)** ZNRF3 (n=6) and **(C)** RSPO-3 (n=10) was determined by qRT-PCR using GAPDH as a housekeeping gene (analysis by paired t-test). **(D)** Protein lysates were prepared and analysed by western blot using an anti-RSPO-3 antibody. Calnexin was used as a loading control (n=7; analysis by unpaired t-test; representative blots shown in the panel above). **(E-G)** HAEC were transfected with RSPO-3 siRNA or scrambled controls and exposed to flow for 48h. Protein lysates were prepared from EC exposed to LMMF and analysed by western blot using **(E)** RSPO-3, **(F)** Frizzled-4 and **(G)** VCAM-1 antibodies. Calnexin was used as a loading control (n=3; analysis by unpaired t-test; representative blots shown in the panels above).

These data suggest that LMMF induces pro-inflammatory changes by increasing the expression of Frizzled-4. Subsequent studies focused on understanding the pathways by which Frizzled-4 determines these and other changes in EC exposed to LMMF.

### Frizzled-4 signalling in EC exposed to LMMF is mediated by increased transcriptional activity of β-catenin

Frizzled-4 is known to activate the canonical, β-catenin-dependent Wnt pathway. We therefore examined whether the anti-inflammatory effects of Frizzled-4 knockdown were associated with altered β-catenin signalling under LMMF conditions. Analysis of fractionated lysates revealed that total β-catenin expression was significantly increased in both cytosolic and nuclear fractions in EC exposed to LMMF relative to HMUF. Nuclear expression of the active, dephosphorylated form of β-catenin was also significantly increased in EC exposed to LMMF (Supplementary Figure S3). Using a reporter assay we demonstrated that LMMF (1h – 48h) significantly increased reporter activity (relative to HMUF), indicating increased β-catenin-mediated transcriptional activity. Reporter activity in EC exposed to LMMF also increased over time (Supplementary Figure S3).

Importantly, reduction of Frizzled-4 expression by siRNA interference resulted in a significant decrease in β-catenin transcriptional activity under LMMF conditions (Figure 3a). This was also associated with reduced β-catenin levels in whole cell lysates (Figure 3b). Moreover, knockdown of RSPO-3 also reduced the expression of β-catenin in whole cell lysates under LMMF conditions (Figure 3b). We therefore investigated whether inhibition of β-catenin transcriptional activity using a small molecule inhibitor, iCRT5^23^, mirrored the effects of Frizzled-4/RSPO-3 knockdown. Addition of iCRT5 to EC for the last 24h of flow exposure significantly inhibited β-catenin transcriptional activity (Figure 3c) and reduced the expression of E-selectin, MCP-1, VCAM-1 and IL-8 transcripts under LMMF conditions relative to vehicle-treated controls (Figure 3d). Similar results were obtained when iCRT5 was added for the duration of flow or when β-catenin expression was reduced following transfection of EC with siRNA targeting β-catenin (Supplementary Figure S4). We also confirmed that iCRT5 significantly reduced the protein expression of VCAM-1 in EC exposed to LMMF (Figure 3e). Furthermore, addition of iCRT5 added for the duration of flow (72h) or for the last 24h, resulted in significantly reduced adhesion of monocytes to EC exposed to LMMF (Figure 3f). These data suggest that Frizzled-4-dependent activation of β-catenin mediates, at least in part, the pro-inflammatory effects of LMMF.

**Figure 3.**
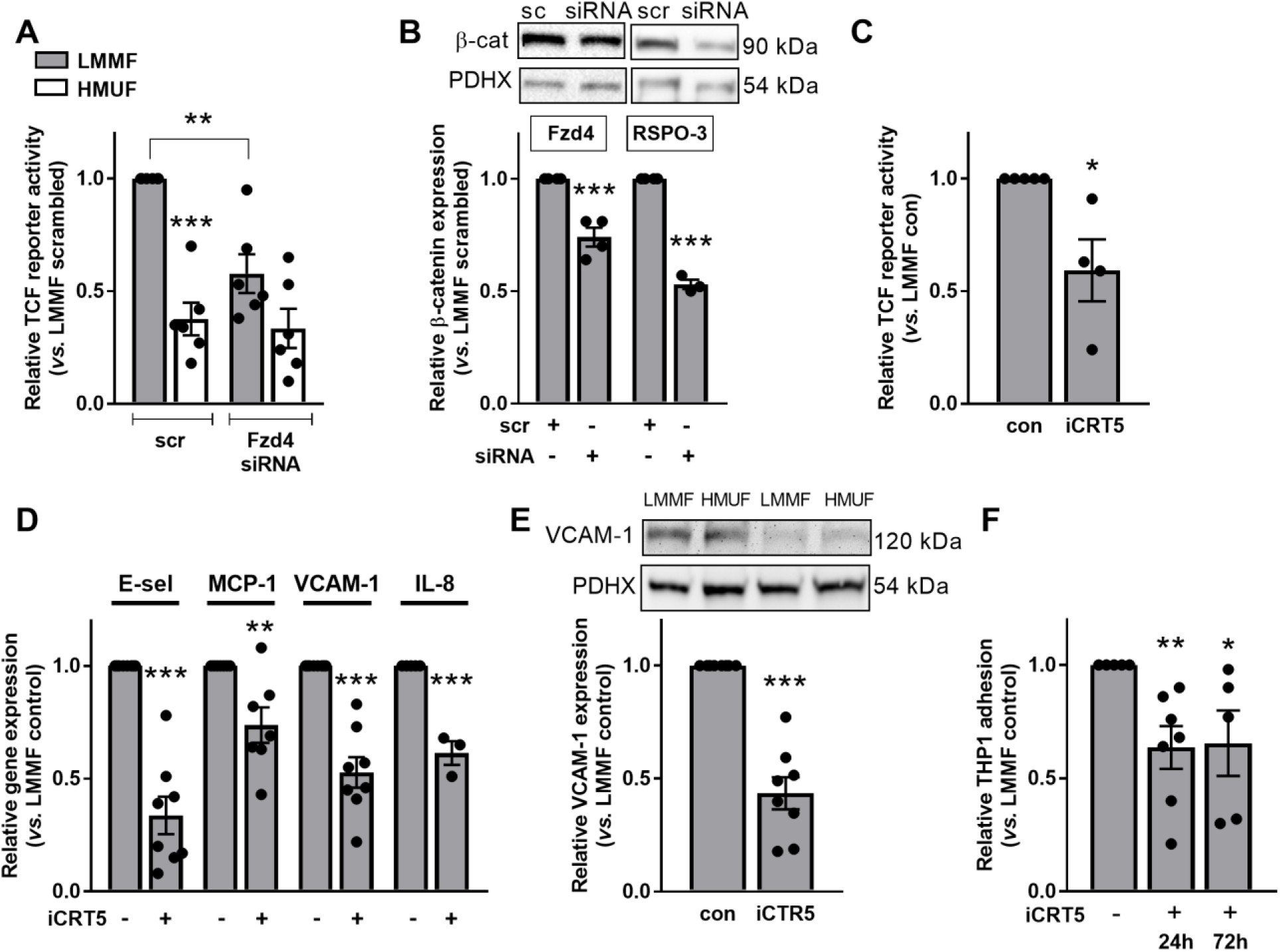
β-catenin activity is increased in EC exposed to LMMF in a Frizzled-4 dependent manner. **(A)** HAEC were transfected with Frizzled-4 siRNA or scrambled control plus TCF reporter constructs and exposed to flow for 48h. Lysates were prepared from cells exposed to LMMF or HMUF and *Firefly* and *Renilla* luciferase activity recorded. Ratios were corrected for protein content of lysates. Results shown relative to LMMF control (n=6; analysis by one-way ANOVA). **(B)** HAEC were transfected with Frizzled-4 or RSPO-3 siRNA or scrambled control and exposed to flow for 48h. Lysates were prepared from EC exposed to LMMF and analysed by western blot using an anti-β-catenin antibody. PDHX was used as a loading control (n=3-4; analysis by unpaired t-test). **(C-F)** HAEC were exposed to flow for 72h and treated with iCRT5 (50μM) for the last 24h of flow exposure. **(C)** HAEC were transfected with TCF reporter constructs 24h prior to flow exposure. Lysates were analysed as in (A). Results shown relative to LMMF control (n=4; analysis by unpaired t-test). **(D)** RNA was harvested from EC exposed to LMMF and gene expression determined by qRT-PCR using GAPDH as a housekeeping gene (n=3-8; analysis by unpaired t-test). **(E)** Lysates were prepared from EC exposed to LMMF and HMUF and analysed by western blot using an anti-VCAM-1 antibody. PDHX was used as a loading control (n=8; analysis by unpaired t-test). **(F)** The number of adherent calcein-labelled THP-1 monocytes was determined in 4 fields of view following treatment with iCRT5 for the final 24h of flow exposure or for the full duration (72h). Results shown relative to untreated controls (n=5-7; analysis by unpaired t-test at each time point.

### Inhibiting β-catenin activity reduces the elevated paracellular permeability seen in monolayers exposed to LMMF

LMMF increases monolayer permeability to FITC-avidin compared to HMUF or static culture; the tracer crosses the endothelium by a paracellular route^24^. Inhibition of β-catenin transcriptional activity using iCRT5 significantly reduced permeability to FITC-avidin under LMMF (Figures 4a and 4b; EC exposed to HMUF included for reference). Closer inspection showed that transport at bicellular junctions rather than multicellular junctions was particularly reduced (Supplementary Figure S5). We subsequently investigated whether inhibiting β-catenin signalling affected the expression of junctional molecules. iCRT5 did not induce changes in the transcript levels of VE-cadherin, ESAM, JAM-B or β-catenin in EC exposed to LMMF. However, it did increase expression of PECAM-1, JAM-A, JAM-C, ZO-1 and Claudin-5 (Supplementary Figure S6). Immunostaining of ZO-1 and VE-cadherin revealed a clear change in junctional organisation following inhibition of β-catenin in EC exposed to LMMF along with increased expression of ZO-1. Junctions appeared disorganised and irregular in control EC exposed to LMMF but more organised following treatment with iCRT5 (Figure 4c-e). The morphology of EC exposed to LMMF was also altered by iCRT5: the cells appeared more consistently oriented and more elongated (representative images in Figure 4c-e). Quantification of the cell shape index confirmed the latter observation (Supplementary Figure S7; HMUF included for reference).

**Figure 4.**
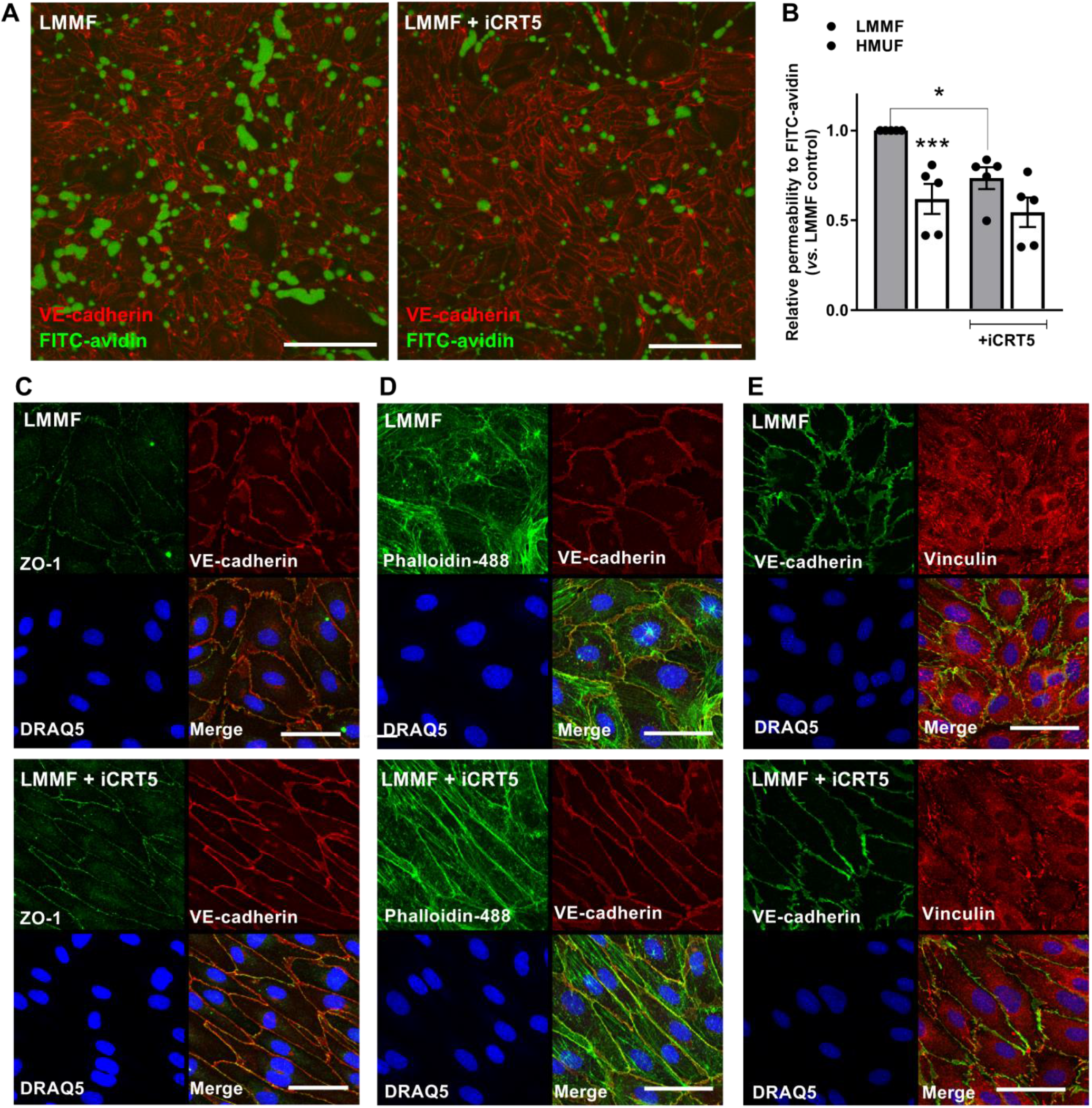
β-catenin activity increases permeability in EC exposed to LMMF. **(A-E)** HAEC were exposed to flow for 72h and treated with iCRT5 (50μM) for the last 24h of flow exposure. **(A)** FITC-avidin was added to monolayers immediately after flow cessation. Images show areas where FITC-avidin binds to biotinylated-gelatin underlying EC. Cells were counterstained with an anti-VE-cadherin antibody and DRAQ5 nuclear stain (images shown are maximum projections of z-stacks; scale = 200μm). **(B)** Accumulation of FITC-avidin was quantified by determining the intensity of FITC-avidin in maximum projections and shown relative to LMMF vehicle control (n=5; analysis by Kruskal-Wallis test with uncorrected Dunn’s test). **(C-E)** EC were fixed and stained with an anti-VE-cadherin antibody and DRAQ5 nuclear stain plus **(C)** anti-ZO-1 antibody **(D)** 488-Phalloidin **(E)** anti-vinculin antibody (scale = 50μm; representative images from 4 independent experiments).

We also assessed the effects of β-catenin inhibition on the actin cytoskeleton in cells exposed to LMMF. Staining with phalloidin revealed striking differences after treatment with iCRT5. There was a reduction in the number of stress fibres and an increase in junctional (cortical) staining that is typically associated with stabilisation of junctions and reduced permeability (Figure 4d)^25^. There was a similar change in cytoskeletal architecture following knockdown of β-catenin or Frizzled-4 (Supplementary Figure S8). The distribution of vinculin under LMMF was also modified by iCRT5: there was greater localisation around junctions compared to the apparent localisation to focal adhesions in untreated cells (Figure 4e). Similar alterations in vinculin localisation were again observed following knockdown of β-catenin or Frizzled-4 (Supplementary Figure S8). Taken together these data suggest that inhibiting β-catenin signalling stabilises cell-cell junctions in EC exposed to LMMF and that this may account for the enhanced barrier properties observed in the presence of iCRT5.

### Inhibition of Wnt5a signalling reduces LMMF-dependent activation of β-catenin and attenuates inflammatory signalling

Since Frizzled receptors are commonly activated by Wnt ligands, we assessed the role of Wnt5a in mediating the response to LMMF. We focused on Wnt5a because it is known to promote inflammatory signalling in EC^26^. We found significantly increased expression of Wnt5a in EC exposed to LMMF compared to HMUF (Figure 5a). Transfection with Wnt5a siRNA resulted in a 50% reduction in Wnt5a expression (Figure 5b) and reduced expression of E-selectin, MCP-1 and VCAM-1 (Figure 5c). Knockdown of Wnt5a also resulted in reduced numbers of stress fibres, increased cortical actin, and apparent elongation of EC exposed to LMMF, similar to the effects observed with knockdown of Frizzled-4 or β-catenin (Figure 5d).

**Figure 5.**
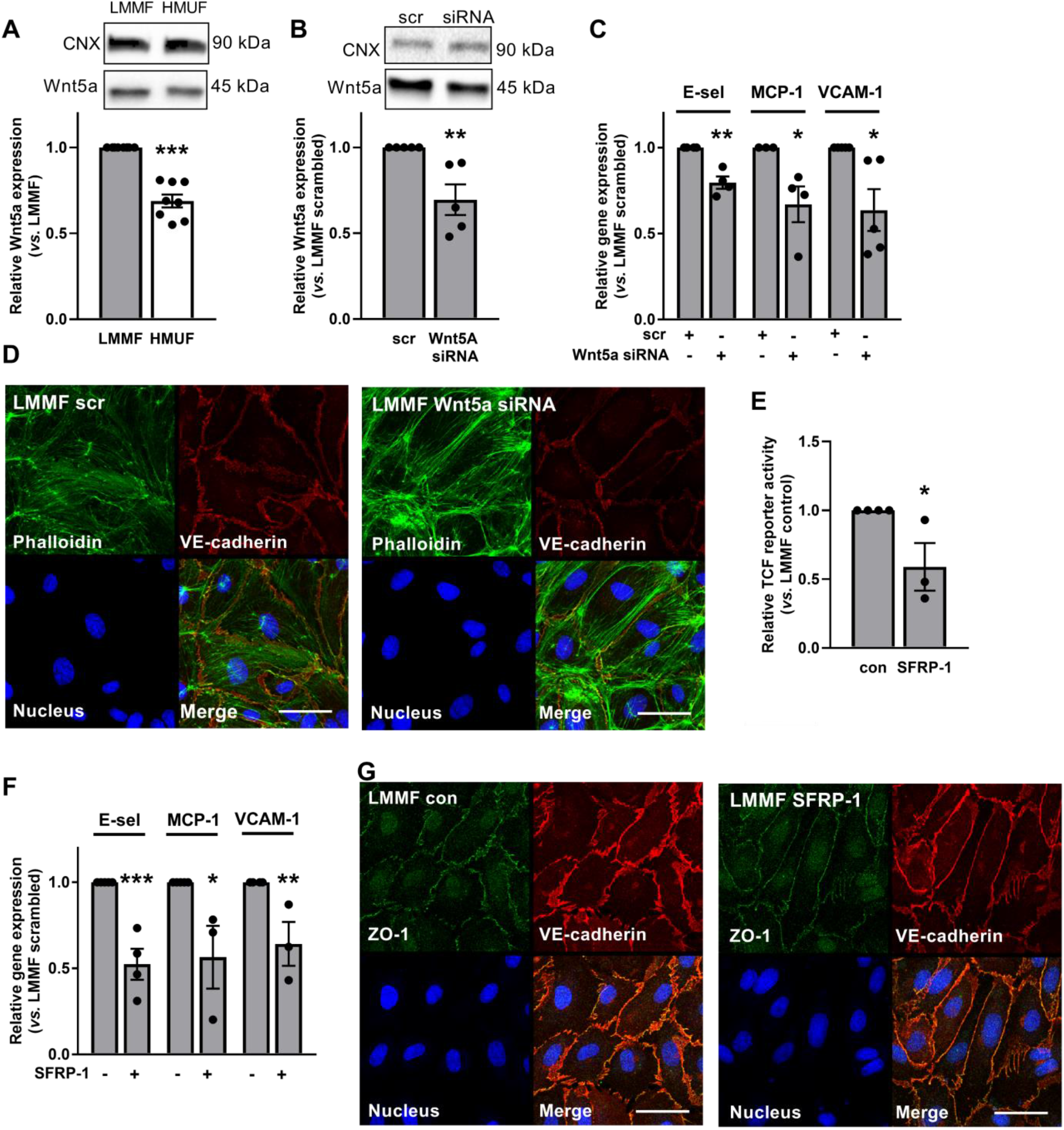
Wnt5a expression is increased in EC exposed to LMMF. **(A)** HAEC were exposed to flow for 72h and the expression of Wnt5a quantified in cells exposed to LMMF or HMUF by western blot using an anti-Wnt5a antibody. Calnexin (CNX) was used as a loading control (n=8; analysis by paired t-test). **(B-D)** HAEC were transfected with Wnt5a siRNA or scrambled controls and exposed to flow for 48h. **(B)** protein and **(C)** RNA was isolated from EC exposed to LMMF. **(B)** Wnt5a expression was assessed by western blot using CNX as a loading control (n=5; analysis by unpaired t-test). **(C)** Gene expression was determined by qRT-PCR using GAPDH as a housekeeping gene (n=4-5; analysis by unpaired t-test). **(D)** EC were fixed and incubated with 488-Phalloidin, anti-VE-cadherin antibody and DRAQ5 nuclear stain and images obtained from 4 fields of view (n=5; representative images shown; scale = 50μm). **(E-G)** HAEC were exposed to flow for 72h and sFRP-1 (200 μg.ml^−1^) added for the final 24h of flow exposure. **(E)** HAEC were transfected with TCF reporter constructs 24h prior to flow exposure. Lysates were prepared from cells exposed to LMMF and *Firefly* and *Renilla* luciferase activity recorded. Ratios were corrected for protein content of lysates. Results shown relative to LMMF control (n=3; analysis by unpaired t-test). **(F)** Gene expression was determined by qRT-PCR using GAPDH as a housekeeping gene (n=3-5; analysis by unpaired t-test). **(G)** EC were fixed and incubated with anti-VE-cadherin and anti-ZO-1 antibodies and DRAQ5 nuclear stain and images obtained from 4 fields of view (n=5; representative images shown; scale = 50μm).

These data suggest that Wnt5a can promote endothelial dysfunction under LMMF conditions. This interpretation was supported by experiments where EC were treated with SFRP-1, which blocks the interaction of Wnt5a with Frizzled receptors^27^ and is shown here to reduce β-catenin reporter activity (Figure 5e). Similar to knockdown of Wnt5a and Frizzled-4 and inhibition of β-catenin signalling, SFRP-1 treatment for 24h significantly reduced the expression of E-selectin, MCP-1 and VCAM-1 in EC exposed to LMMF (Supplementary Figure S9). Interestingly, inhibition was greater when cells were exposed to SFRP1 for 72h (Figure 5f). Furthermore, SFRP-1 also resulted in junctional and cytoskeletal changes similar to those observed with iCRT5 (Figure 5g and Supplementary Figure S9).

### Frizzled-4 signalling in EC exposed to LMMF is independent of Lrp5/6 but may require Ryk signalling

The data presented so far suggests that EC dysfunction under LMMF conditions can be promoted via Wnt5a-Frizzled-4 signalling. The requirement for β-catenin is consistent with the involvement of a canonical (β-catenin-dependent) Wnt signalling pathway, and this is supported by the finding that phosphorylation of GSK3β(Ser9) was increased in EC by exposure to LMMF rather than HMUF (Figure 6a). Phosphorylation of the Ser9 residue inhibits GSK3β and is a necessary step in the activation of β-catenin; it results in inhibition of the β-catenin destruction complex leading to the nuclear localisation of dephosphorylated (active) β-catenin. GSK3β phosphorylation on Ser9 was significantly reduced following knockdown of Frizzled-4 or RSPO-3 (Figure 6b) suggesting a role for the destruction complex in mediating their effects on β-catenin signalling in EC exposed to LMMF. We consequently investigated whether stabilisation of the β-catenin destruction complex affected the activation of β-catenin and the phenotype of EC exposed to LMMF. We found that treating the cells with IWR-1 (which stabilises Axin2 levels) inhibited the transcriptional activity of β-catenin and reduced the expression of E-selectin and MCP-1 under LMMF conditions, mimicking the effects of Wnt5a and Frizzled-4 knockdown and inhibition of β-catenin signalling (Figure 6c-d). Stabilisation of the destruction complex with IWR-1 also resulted in morphological, junctional and cytoskeletal changes similar to those observed previously (Figure 6e and Supplementary Figure S7). These data are consistent with a role for activation of a Frizzled-4-dependent canonical Wnt pathway in cells exposed to LMMF.

**Figure 6.**
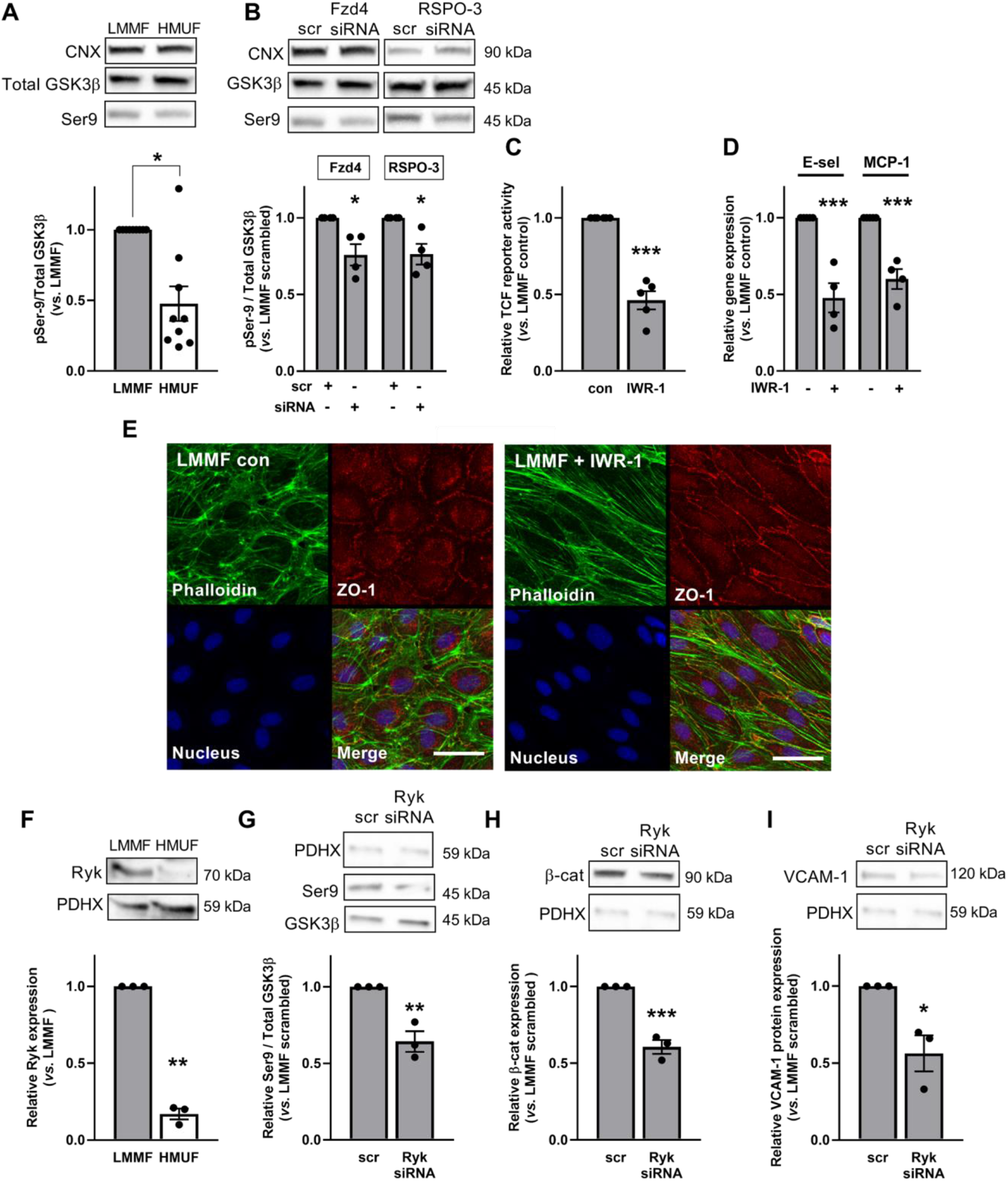
LMMF inhibits the β-catenin destruction complex in a Frizzled-4 and RSPO-3 and Ryk-dependent manner. **(A-B)** GSK3β phosphorylation was quantified by western blot using anti-phospho Ser9 and anti-GSK3β (total) antibodies in protein lysates from HAEC exposed to **(A)** LMMF and HMUF for 72h or **(B)** LMMF for 48h following transfection with Frizzled-4 or RSPO-3 siRNA or scrambled controls. Calnexin (CNX) was used as a loading control. **(A)** Results shown relative to LMMF control (n=9; analysis by Wilcoxon signed rank test). **(B)** Results shown relative to LMMF scrambled (n=4; analysis by unpaired t-test). **(C-E)** HAEC were exposed to flow for 72h and IWR-1 (10μM) added for the final 24h of flow exposure. **(C)** HAEC were transfected with TCF reporter constructs 24h prior to flow exposure. Lysates were prepared from cells exposed to LMMF and *Firefly* and *Renilla* luciferase activity recorded. Ratios were corrected for protein content of lysates. Results shown relative to LMMF control (n=5; analysis by unpaired t-test). **(D)** Gene expression was determined by qRT-PCR using GAPDH as a housekeeping gene (n=4; analysis by unpaired t-test). **(E)** EC were fixed and stained with 488-Phalloidin, anti-VE-cadherin antibody and DRAQ5 nuclear stain. Images obtained from 4 fields of view (n=5; representative images shown; scale = 50μm). **(F)** Protein lysates were obtained from HAEC exposed to LMMF and HMUF for 72h or **(G-I)** from HAEC exposed to LMMF for 48h following transfection with Ryk siRNA or scrambled controls. **(F-I)** Expression of proteins was determined by western blot using antibodies targeting **(F)** Ryk, **(G)** phosphor-Ser9 and anti-GSK3β, **(H)** β-catenin and **(I)** VCAM-1. PDHX was used as a loading control (representative blots shown in the panels above; n=3; analysis by unpaired t-test).

To test this further, we assessed the activation of Lrp6 (a hallmark of the canonical Wnt pathway) under flow conditions. Surprisingly, we found no evidence of more Lrp6 phosphorylation under LMMF than under HMUF (Supplementary Figure S10). Moreover, inhibiting canonical Wnt signalling by treating EC with DKK-1 (which blocks the interaction of Lrp and Frizzled receptors) for 24h did not have the expected effect on the expression of E-selectin or MCP-1, THP-1 monocyte adhesion, cell-shape index or cytoskeletal/junctional organisation (Supplementary Figure S7 and S9).

Because these data suggest that Frizzled-4 activates β-catenin independently of Lrp signalling, we investigated alternative Wnt pathway components that could mediate the effects of Frizzled-4 activation and found that Ryk expression was significantly increased in EC exposed to LMMF for 72h (Figure 6f). Knockdown of Ryk was associated with reduced phosphorylation of GSK3β on Ser9 (Figure 6g) and reduced levels of β-catenin (Figure 6h), consistent with a role in regulating β-catenin under LMMF conditions. Moreover, knockdown of Ryk reduced the expression of VCAM-1 in cells exposed to LMMF (Figure 6i).

## Discussion

Here we provide evidence of a novel Frizzled-4-dependent signalling pathway that is activated in EC in response to LMMF. The expression of Frizzled-4 is significantly higher under LMMF compared to HMUF, which is dependent on increased expression of RSPO-3. In response to LMMF, Frizzled-4 increases the transcriptional activation of β-catenin which occurs independently of Lrp signalling. β-catenin promotes inflammatory activation and barrier disruption associated with LMMF (summarised in the Figure 7).

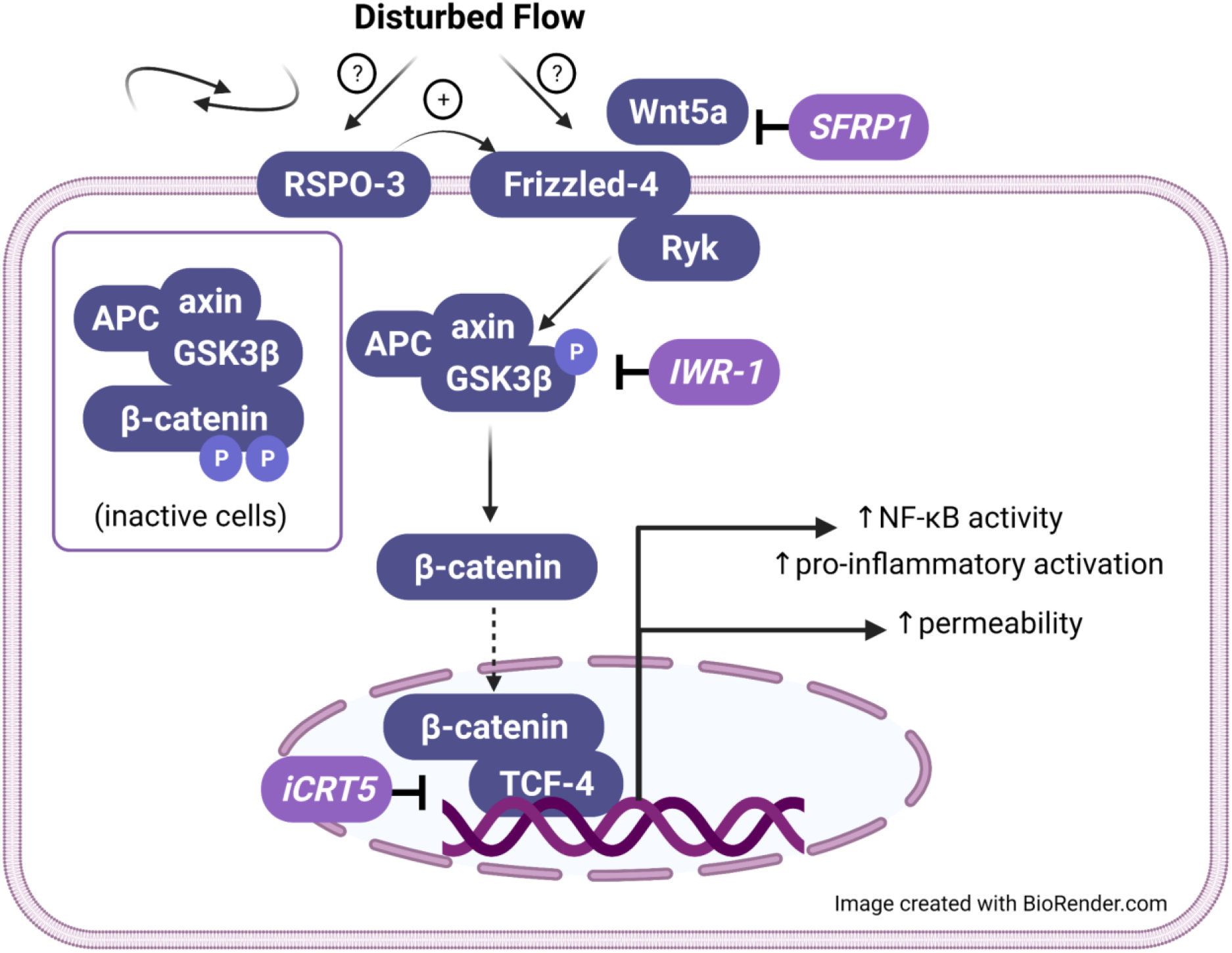

To our knowledge this is the first evidence that Frizzled-4 protein expression is elevated in response to LMMF. Frizzled-4 transcript levels decreased under the same conditions. This observation is in line with data obtained following surgical induction of flow disturbance in mouse carotid arteries^28^, but a second transcriptomic analysis found *increased* Frizzled-4 transcript levels in the inner curvature of the porcine aortic arch; this may reflect an adaptation to chronic LMMF exposure ^7^. Regardless of the apparent discrepancies in transcript levels between earlier studies, we found increased Frizzled-4 protein in the inner curvature of the mouse aortic arch, which is chronically exposed to LMMF, suggesting that the increased protein expression in our cell culture model is physiologically relevant.

Since Frizzled-4 protein levels were elevated under LMMF without any concomitant rise in transcript levels, we investigated other mechanisms of regulation. E3 ubiquitin ligases ZNRF3 and RNF43 target Frizzled for proteasomal degradation and thus negatively regulate Frizzled signalling^29^. We did not observe any change in expression of these enzymes under flow conditions. However, we did find a significant increase in the expression of RSPO-3, which inhibits the activity of ZNRF3^29^, in EC exposed to LMMF. This plausibly explains the increase in Frizzled-4. Knockdown of RSPO-3 significantly lowered the expression of Frizzled-4 in cells exposed to LMMF and reduced the inflammatory activation induced by this type of flow further supporting a role for RSPO-3 in the regulation of Frizzled-4 under LMMF.

Whilst we have shown that reducing Frizzled-4 attenuates LMMF-induced inflammatory activation, the mechanisms by which Frizzled-4 signalling is activated are not fully defined. One possibility is that, as a G-protein coupled receptor, Frizzled-4 may be directly activated by mechanical force, as has been demonstrated for other GCPRs^30^. Another is Wnt-mediated activation. Our data support a role for Wnt5a in mediating LMMF-induced inflammatory activation: the expression of Wnt5a was increased in EC exposed to LMMF and knockdown of Wnt5a reduced the expression of proinflammatory genes under LMMF. Wnt5a has previously been shown to promote endothelial dysfunction in static EC ^26,31^ via a JNK-dependent pathway^26^, although we found no evidence of JNK activation in cells exposed to LMMF (data not shown).

Historically, it was thought that Wnt5a promotes inflammatory signalling by activating noncanonical (i.e. β-catenin-independent) pathways. However, Wnt5a can bind to and promote Frizzled-4 internalisation^32^ and oligomerisation^33^ and, in the presence of Frizzled-4, can activate canonical Wnt signalling^34^. Our data add to evidence for the importance of ligand-mediated activation of Frizzled-4 by showing that LMMF-induced activation of β-catenin and proinflammatory signalling is also reduced following treatment with SFRP-1, which binds and antagonises Frizzled-4^27^, although SFRP-1 may have actions at other Frizzled receptors^27,35^.

Knockdown of Frizzled-4 and RSPO-3 reduced the activation of β-catenin in EC exposed to LMMF. This supports a role for a canonical Wnt pathway activated under such flow. Our observation of increased GSK3β also indicates the presence of a flow-sensitive canonical pathway. However, we could not detect the phosphorylation of Lrp6 that is typically associated with canonical Wnt signalling. Furthermore, inhibition of the canonical pathway using DKK-1 (which blocks the interaction between Frizzled and Lrp co-receptors) had no effect on EC responses to LMMF. It is possible that upon stimulation, Frizzled-4 directly activates the β-catenin destruction complex, independently of Lrp5/6. This could occur as a result of dimerization of Frizzled-4 receptors and consequent clustering of Dishevelled (Dvl) proteins^36^, leading to the dimerization and polymerisation that is necessary for activation of the axin degradasome^37^. Alternatively, Wnt5a-Frizzled-4 signalling may occur independently of Dvl, as has been demonstrated during neurite outgrowth^38^. Intriguingly, it has also been documented that Wnt5a can promote endothelial dysfunction via the Wnt co-receptor, Ryk, an atypical receptor kinase that lacks kinase activity^39^. Ryk can associate with Frizzled receptors and Dvl in 293T cells and this is required for transcriptional activity of TCF4^40^. However, a direct interaction with Wnt pathway components has not previously been demonstrated in EC. We demonstrated here that Ryk expression is increased in EC exposed to LMMF and that depletion of Ryk reduces GSK3β phosphorylation and VCAM-1 expression under LMMF. It is therefore possible that Ryk is recruited to Frizzled-4 receptors upon ligation by Wnt5a and contributes to the activation of β-catenin. Taken together, our data suggest the presence of an atypical Wnt signalling pathway that is activated by LMMF and promotes endothelial dysfunction. Interestingly, Gelfand et.al also provide evidence that PECAM-1 plays a role in the regulation of β-catenin under atherogenic flow conditions^41^ and that this could be due to phosphorylation and consequent inhibition of GSK3β^42^; LMMF-induced activation of β-catenin is complex and may require multiple converging pathways.

We have shown here that the increased expression of RSPO-3 and Frizzled-4 in EC exposed to LMMF results in increased nuclear translocation and transcriptional activation of β-catenin. Similar findings were previously obtained in EC exposed to low and oscillatory shear stress for 6-24h^41,43^, suggesting that activation of β-catenin is common under atherogenic flow conditions. We also demonstrated that increased transcriptional activity of β-catenin resulted in increased expression of pro-inflammatory molecules (VCAM-1, IL-8, MCP-1, E-selectin) and increased adhesion of THP-1 monocytes in EC exposed to LMMF.

In addition to its actions in proinflammatory signalling, our data reveal important effects of flow-activated Wnt signalling and β-catenin transcriptional activity on cell morphology, cytoskeletal organisation and permeability. Ghim et al recently demonstrated that monolayers exposed to LMMF exhibit increased permeability to FITC-avidin, which we confirmed here^20^. We additionally found that inhibiting β-catenin transcriptional activity with a small molecule inhibitor significantly reduced the permeability. This predominantly reflected effects on bicellular junctions.

EC exposed to LMMF exhibited a randomly oriented cobblestone morphology and a disorganised actin cytoskeleton with numerous stress fibres. Immunostaining for ZO-1, PECAM-1 and VE-cadherin revealed irregular and disorganised junctions. Inhibition of the flow-dependent Wnt pathway at various levels appeared to promote the formation of more organised junctions, plausibly accounting for the reduction in permeability it caused. Although expression of junctional proteins did not appear to be altered, we did observe a significant increase in JAM-A, JAM-C, Claudin-5 and ZO-1 transcript levels following inhibition of β-catenin transcriptional activity A role for ZO-1 in maintaining VE-cadherin function has been seen in EC cultured under static conditions: ZO-1 increases tensile force on VE-cadherin, promotes the formation of cortical actomyosin structures and recruits vinculin to adherens junctions^44^. We also demonstrated in this study that blockade of the LMMF-induced Wnt pathway is associated with redistribution of vinculin to cell-cell contacts. This is consistent with other evidence suggesting that vinculin binds to VE-cadherin and stabilises adherens junctions during force-dependent remodelling^45^.

Blockade of the Wnt pathway and β-catenin transcriptional activity was also associated with EC elongation and alignment, accompanied by the formation of the dense cortical actin ring typically observed in EC exposed to unidirectional shear stress^25,46^ and associated with enhanced barrier function^47^. Recent studies provide evidence that EC re-orient and align themselves so as to minimise transverse wall shear stress^48^ and that EC fail to remodel when transverse wall shear stress is high, resulting in inflammatory activation^49^ and increased permeability^24^. It is known that unidirectional shear stress induces cytoskeletal remodelling and consequent permeability reduction via rapid activation and translocation of Rac and cortactin to cell-cell junctions^46^, along with paxillin and FAK^50^. However, the mechanisms governing permeability and cytoskeletal organisation under disturbed flow are less well defined. Here we report that following inhibition of the flow-dependent Wnt pathway at various levels, EC appear to undergo dramatic remodelling of junctions and the cytoskeleton even though transverse wall shear stress remains high. Several Wnt pathway components can interact with and regulate the cytoskeleton^51^, but our data point towards transcriptional regulation of cytoskeletal components or effectors. It is also possible that blocking Frizzled-4/β-catenin signalling interferes with a directional flow sensor so that cells are less able to respond to LMMF.

Our finding that β-catenin-dependent transcriptional activity alters cytoskeletal organisation in response to LMMF via Wnt5a-Frizzled-4 signalling is supported by a previous study of human coronary artery EC cultured under static conditions. Here, gene expression profiling revealed enrichment of genes involved in cytoskeletal remodelling following exposure to Wnt5a, which was associated with increased endothelial permeability^39^. The effects of Wnt5a were mediated by Ryk^39^, which is consistent with our data showing a potential role for Ryk in mediating responses to LMMF. Although the authors did not explore whether Frizzled-4 and β-catenin were involved, a subsequent study found that RSPO-3 was also associated with barrier disruption in EC cultured under static conditions; in this case, downstream mechanisms were not studied^52^. We speculate that under LMMF, elevated RSPO-3 and Wnt5a act synergistically to increase signalling through Frizzled-4 and Ryk, leading to increased transcriptional activity of β-catenin and expression of genes that promote the formation of stress fibres, disorganisation of adherens junctions, increased permeability and endothelial dysfunction.

In conclusion, our results demonstrate that inhibition or deletion of β-catenin reduces inflammatory signalling and enhances barrier function thus pointing towards a role for β-catenin in LMMF-induced endothelial dysfunction. However, our previous research demonstrates that β-catenin can also play a pro-survival role in EC exposed to LMMF^21^. We have also shown that β-catenin can interact with eNOS in EC^53^ and that β-catenin is required for maximal activation of eNOS in EC exposed to HMUF^21^. The functions of β-catenin under flow conditions are thus complex. Further research is required to understand the dual functions of β-catenin signalling in EC, and further understanding of upstream and downstream pathways is necessary before such signalling can form the basis of therapeutic interventions.

## Acknowledgements

This work was supported by funding from The British Heart Foundation [FS/16/2/31739] to CW and PG/15/102/31890 to PDW

