## Supplementary information for "Frizzled-4 regulates β-catenin in endothelial cells exposed to disturbed flow via an atypical Wnt pathway leading to proinflammatory activation and increased permeability"

### SUPPLEMENTARY FIGURES

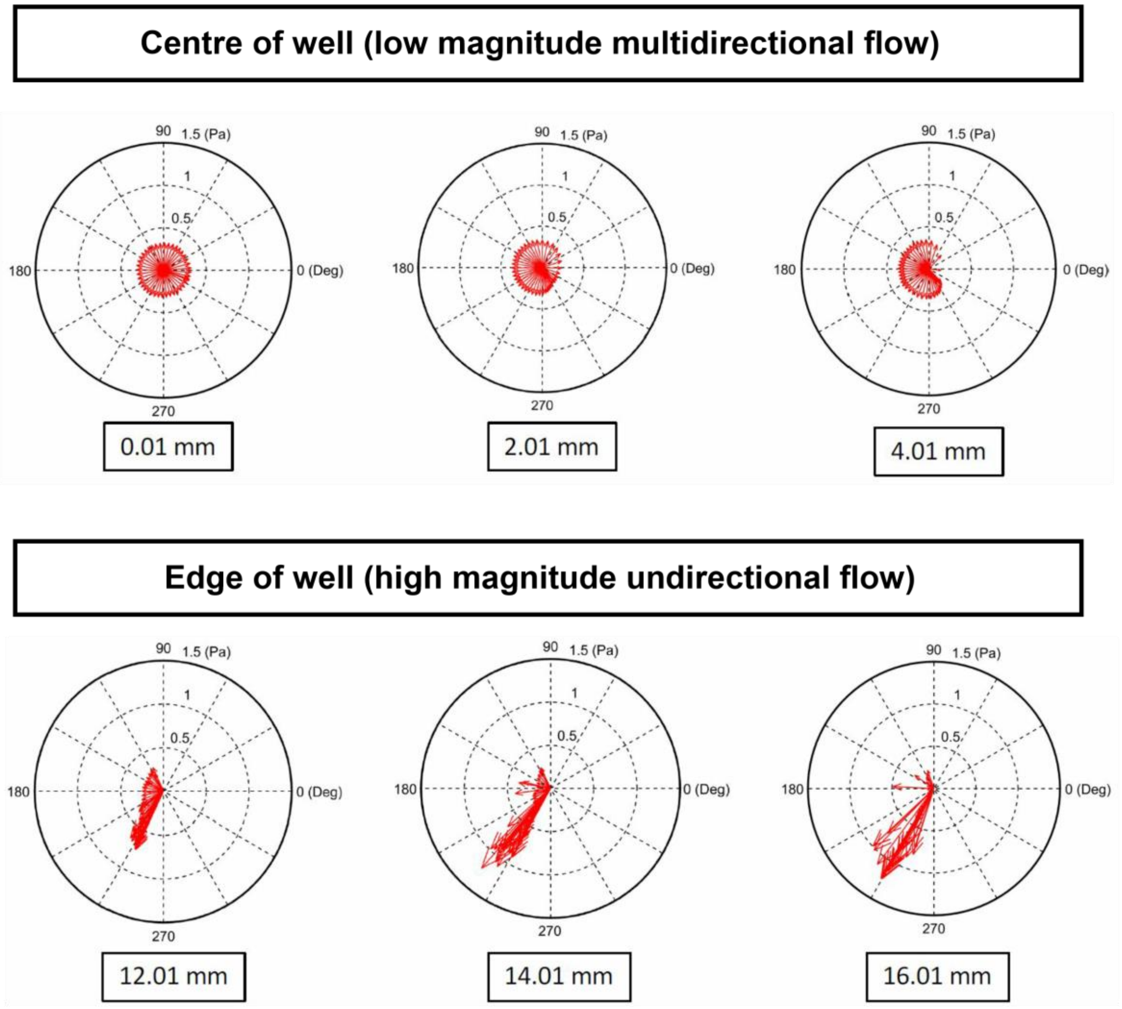


**Supplementary Figure 1. Polar plots showing direction of WSS vectors in swirling wells**

Individual polar plots of the magnitude and direction of instantaneous WSS vectors during one cycle. Each plot applies to one radial distance from the centre of the well. Each arrow represents an instantaneous WSS vector, whose length represents its magnitude, which is plotted at angles corresponding to the shear direction.

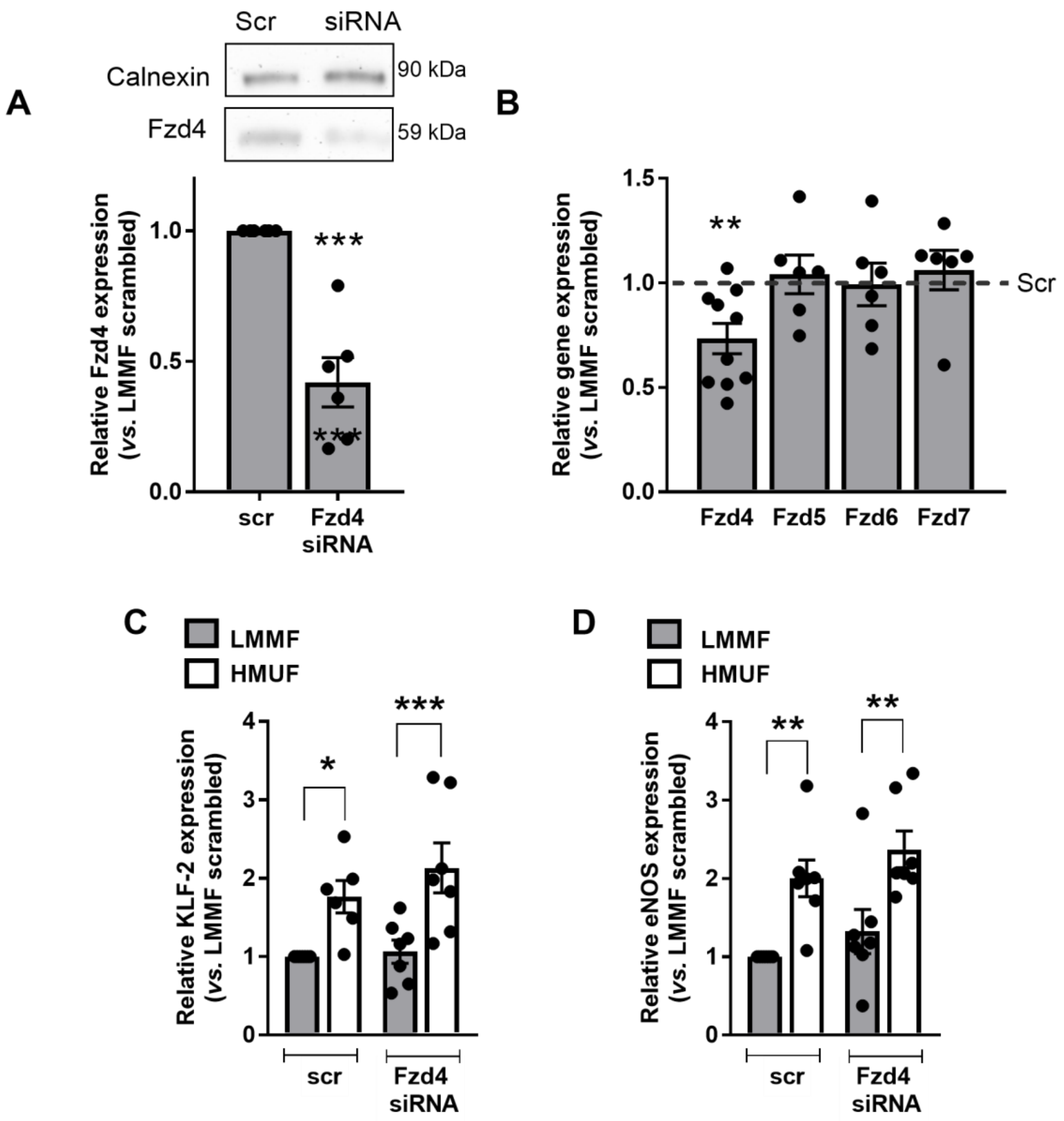


**Supplementary Figure 2. Efficacy and specificity of Frizzled-4 knockdown using siRNA**

**(A-H)** HAEC were exposed to flow for 48h following transfection with Frizzled-4 siRNA or scrambled controls. **(A)** Protein lysates were obtained from HAEC exposed to LMMF and the expression of Frizzled-4 assessed by western blot using calnexin as a loading control (n=6; analysis by unpaired t-test). **(B)** RNA was harvested from EC exposed to LMMF and the expression of Frizzled-4, Frizzled-5, Frizzled-6 and Frizzled-7 was determined by qRT-PCR using GAPDH as a housekeeping gene (n=6-10; analysis by unpaired t-test; results shown relative to LMMF scrambled control (dashed line)). **(C-D)** RNA was harvested from EC exposed to LMMF and HMUF and the expression of **(C)** KLF-2 and **(D)** eNOS was determined by qRT-PCR using GAPDH as a housekeeping gene (n=6-7; analysis by Kruskal-Wallis test with uncorrected Dunn's test; results shown relative to LMMF scrambled control).

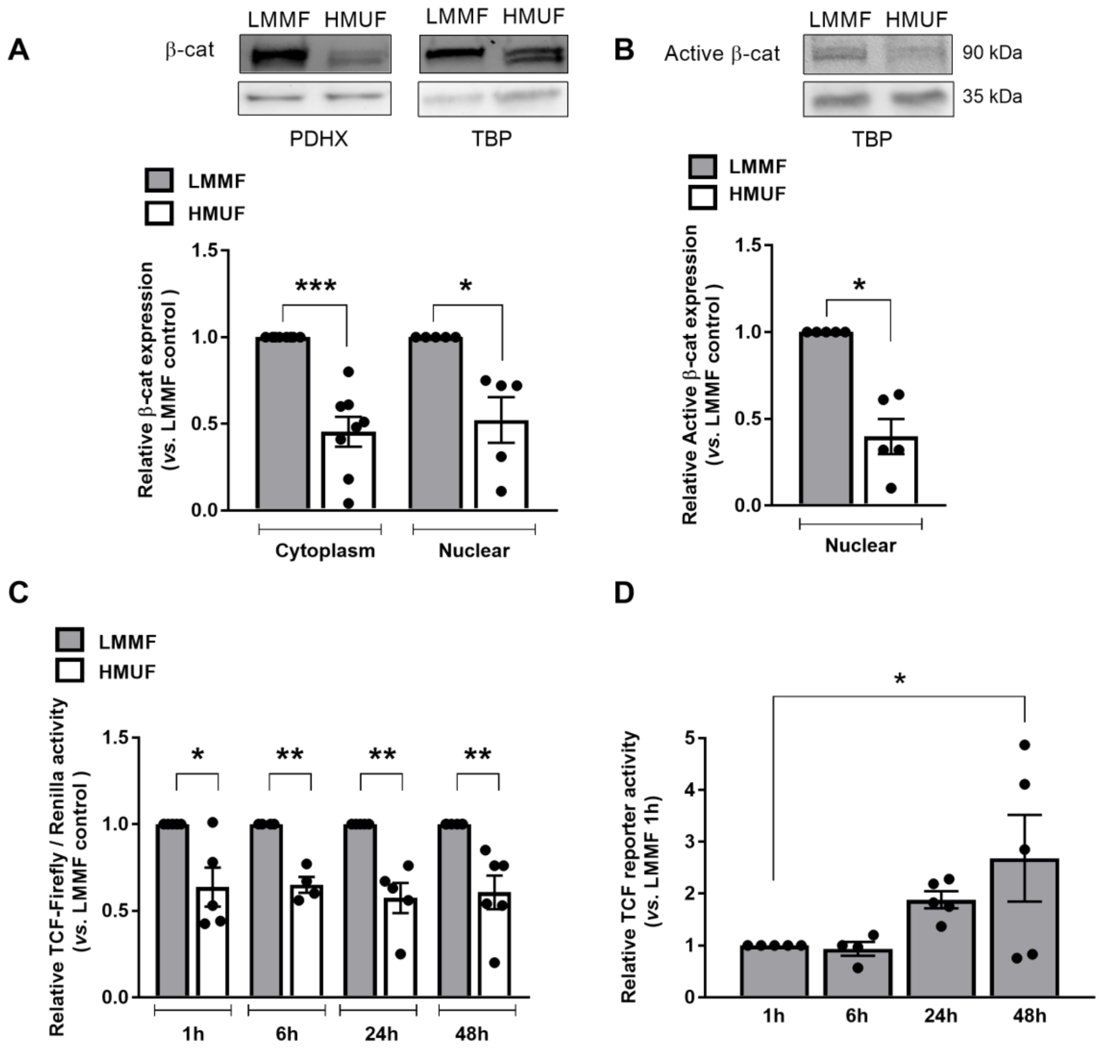


**Supplementary Figure 3. LMMF increases the expression and activation of β-catenin in human aortic endothelial cells**

**(A-B)** HAEC were exposed to flow 72h and lysates prepared from EC exposed to LMMF and HMUF subject to sub-cellular fractionation. Cytosolic and nuclear fractions were analysed by western blot using **(A)** total β-catenin (n=8) and **(B)** active β-catenin antibodies (n=5). PDHX and TBP were used as loading controls for cytosolic and nuclear fractions respectively; analysis by paired t-test; representative blots shown in the panels above. **(C)** HAEC were transfected with Cignal TCF/LEF reporter construct prior to flow exposure for 1h – 48h. Lysates were prepared from cells exposed to LMMF or HMUF and *Firefly* and *Renilla* luciferase activity was recorded. Ratios were corrected for protein content of lysates. **(C)** Results shown relative to LMMF control at each time point and analysed by paired t-test at each time point (n=4-6). **(D)** Results shown relative to LMMF 1h and analysed by one-way ANOVA (n=4-6).

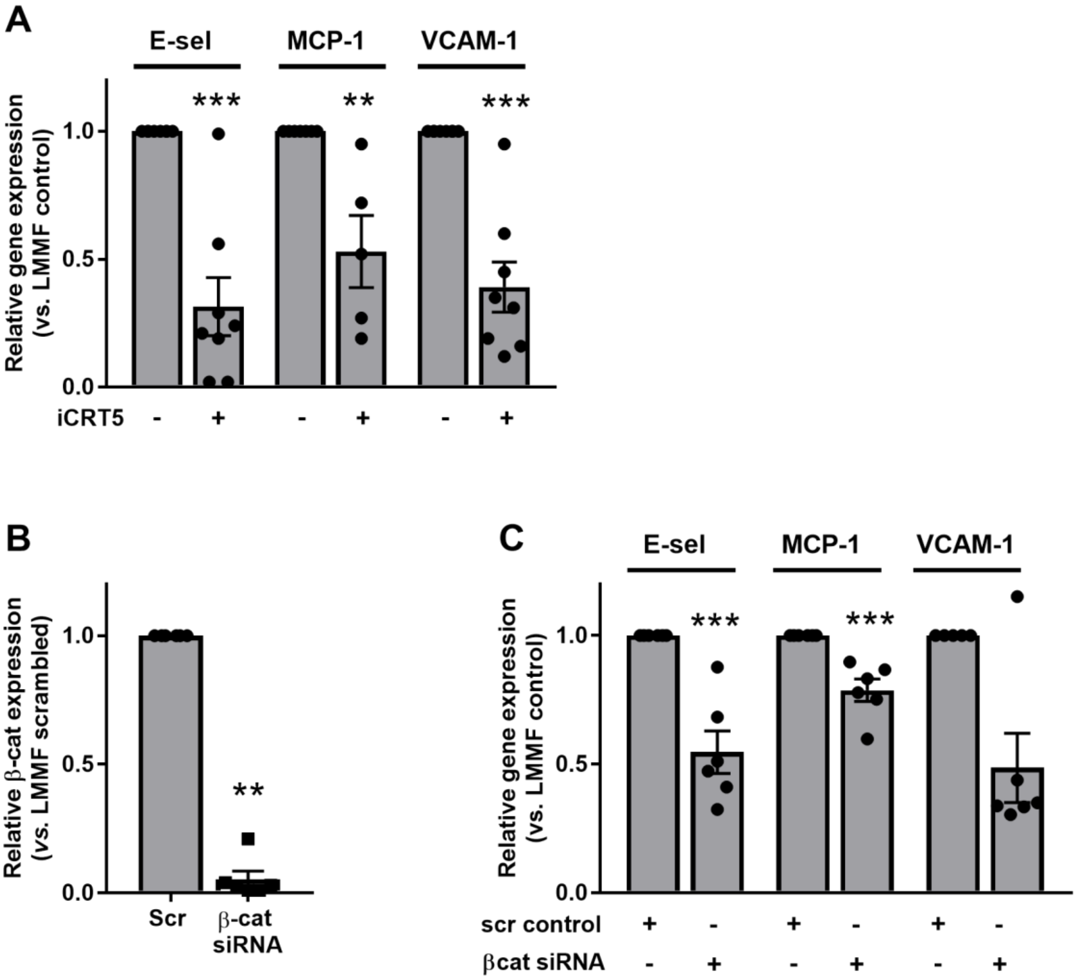


**Supplementary Figure 4. Expression of pro-inflammatory genes in EC exposed to LMMF following inhibition or knockdown of β-catenin**

**(A)** HAEC were exposed to flow for 72h with iCRT5 (50 µM) added for the final 24h of flow exposure. RNA was harvested from EC exposed to LMMF or HMUF and the expression of E-selectin, MCP-1 and VCAM-1 determined by qRT-PCR using GAPDH as a housekeeping gene (n=5-8; analysis by unpaired t-test). **(B-C)** HAEC were exposed to flow for 72h following transfection with β-catenin siRNA. **(B)** The expression of β-catenin protein was assessed by western blot using PDHX as a loading control (n=6; analysis by Mann-Whitney test; representative images shown in panel above). **(C)** RNA was harvested from EC exposed to LMMF and the expression of pro-inflammatory genes assessed by qRT-PCR using GAPDH as a housekeeping gene (n=6; analysis by Mann-Whitney test).

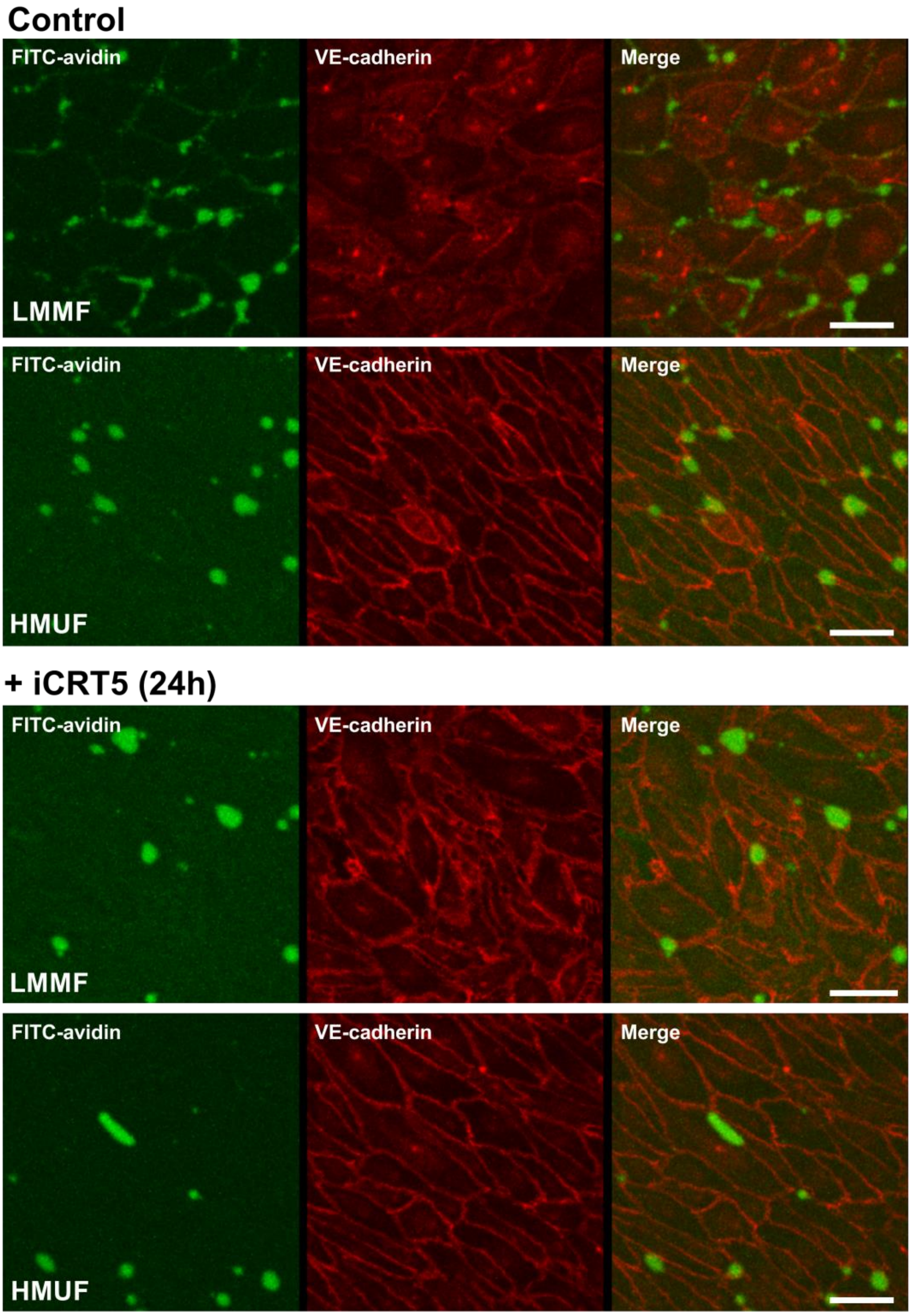


**Supplementary Figure 5. Permeability of HAEC in the presence of iCRT5**

HAEC were cultured on biotinylated-gelatin and exposed to flow for 72h then treated with iCRT5 (50µM) for the last 24h of flow exposure. FITC-avidin was added to monolayers immediately after flow cessation. Images show areas where FITC-avidin binds to biotinylated-gelatin underlying EC. Cells were counterstained with an anti-VE-cadherin antibody and DRAQ5 nuclear stain (images shown are maximum projections of z-stacks; scale = 50µm)

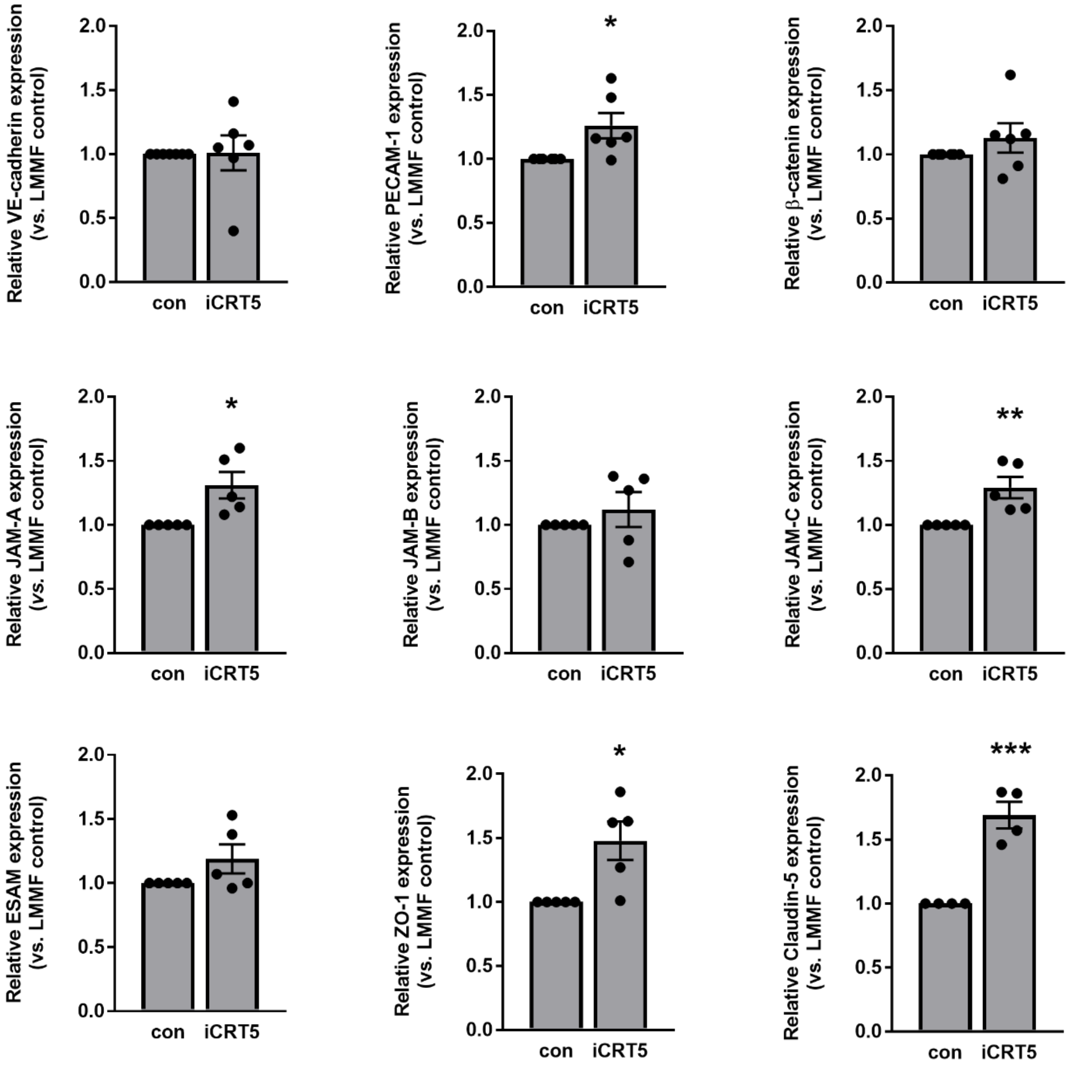


**Supplementary Figure 6. Transcript levels of junctional proteins following inhibition of β-catenin signalling**

HAEC were exposed to flow for 72h and treated with iCRT5 (50µM) for the last 24h of flow exposure. RNA was harvested from EC exposed to LMMF and expression of VE-cadherin, PECAM-1, β-catenin, JAM-A, JAM-B, JAM-C, ESAM**,** ZO-1 and Claudin-5 assessed by qRT-PCR using GAPDH as a housekeeping gene (n=5-6; analysis by unpaired t-test).

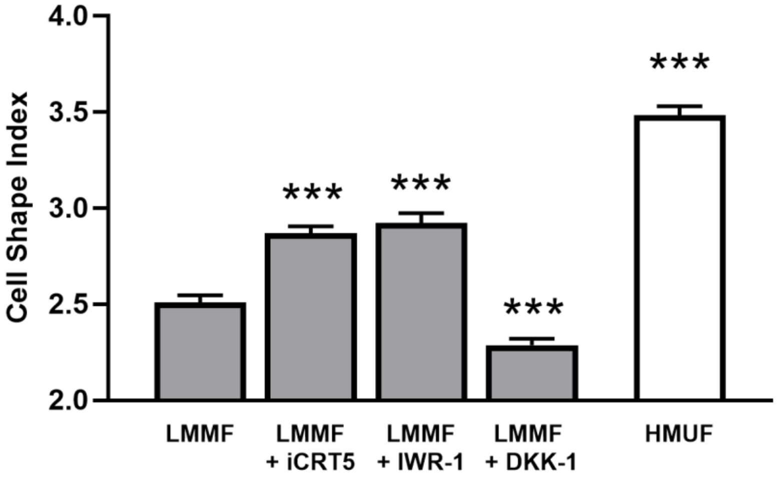


**Supplementary Figure 7. Quantification of endothelial morphology following inhibition of Wnt-β-catenin signalling**

HAEC were fixed and stained with VE-cadherin and DRAQ5 and viewed at x10 magnification. The shape index of all cells in at least 3 fields of view per experiment (approx. 200 cells per field) was determined in cells exposed to LMMF and treated with iCRT5 (50 µM), IWR-1 (10 µM) or DKK-1 (250 ng.ml$^{-1}$). Shape indices of EC exposed to HMUF are shown for comparison (n=3; analysis by Kruskal-Wallis test with uncorrected Dunn's test).

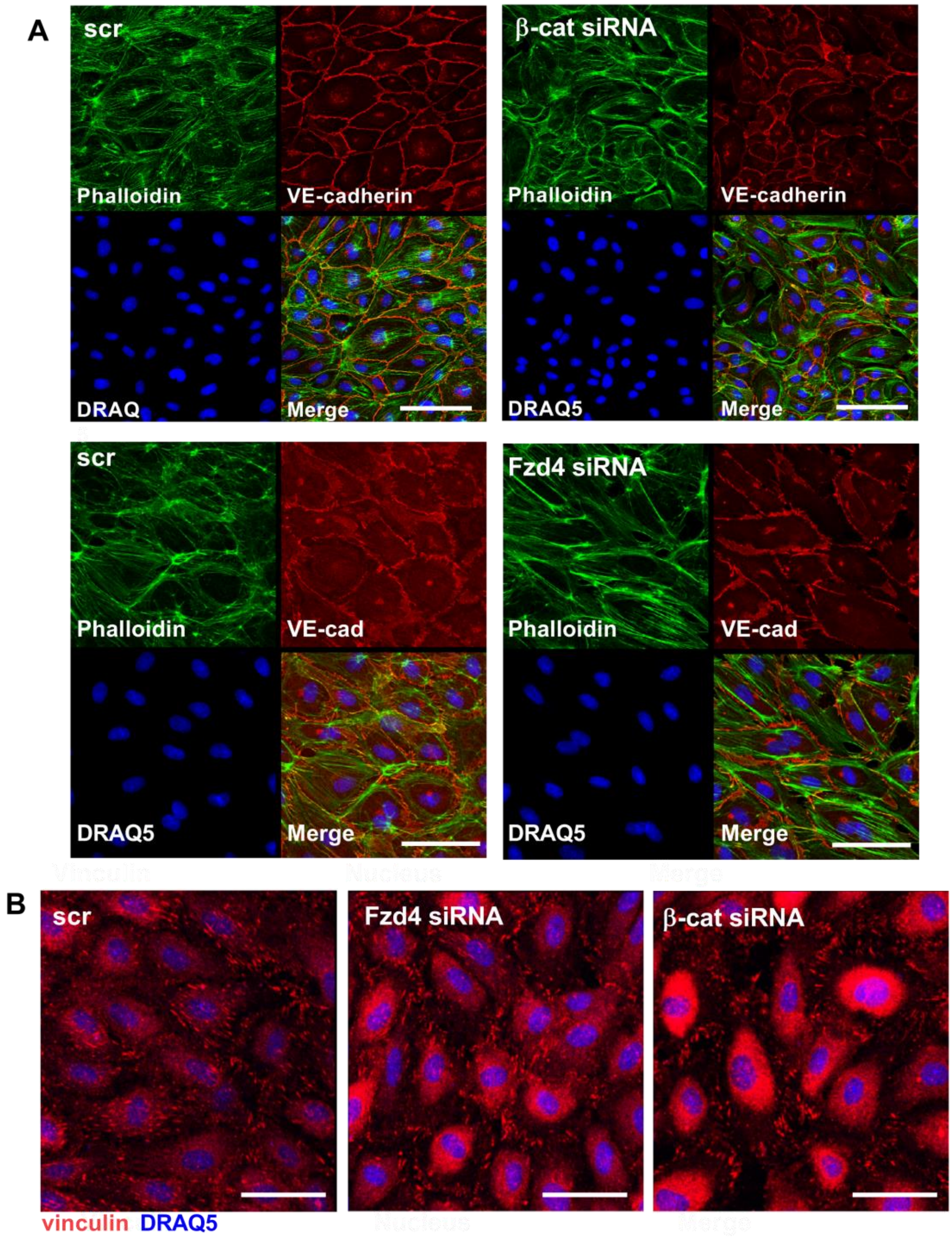


**Supplementary Figure 8. Knockdown of β-catenin or Frizzled-4 alters organisation of the cytoskeleton and vinculin in EC exposed to LMMF**

**(A)** HAEC were exposed to flow for 48h following transfection with siRNA targeting Frizzled-4 or β-catenin and compared to scrambled-transfected control. EC were fixed and incubated with anti-VE-cadherin antibody, 488-Phalloidin and DRAQ5 nuclear stain (n=5; representative images shown). scale = 100 µm for β-catenin siRNA and 50 µm for Frizzled-4 siRNA. **(B)** HAEC were exposed to flow for 72h following transfection with siRNA targeting Frizzled-4 or β-catenin and compared to scrambled-transfected control. EC were fixed and incubated with anti-vinculin antibody and DRAQ5 nuclear stain (n=4; scale = 50 µm; representative images shown).

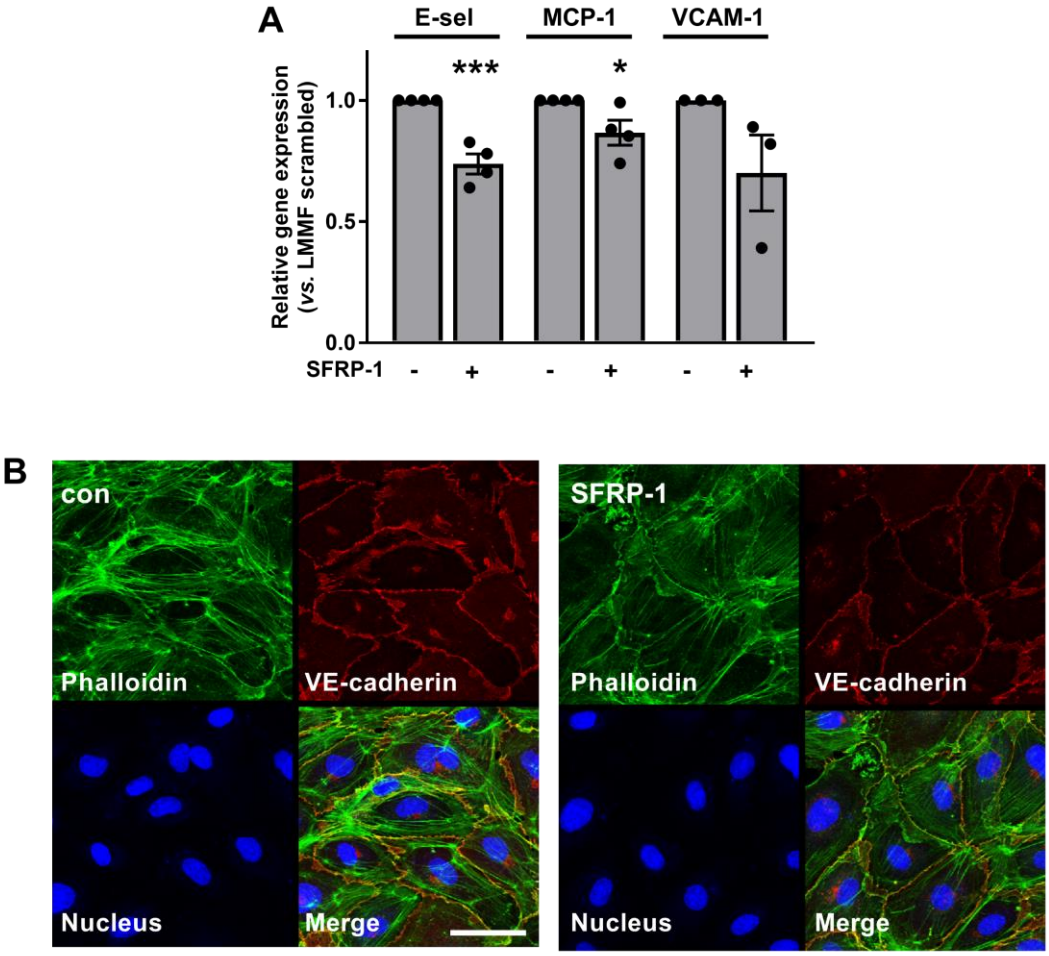


**Supplementary Figure 9. SFRP-1 reduces inflammatory signalling in EC exposed to LMMF and alters cytoskeletal organisation**

**(A-B)** HAEC were exposed to flow for 72h and treated with SFRP-1 (100 ng.ml$^{-1}$) for the last 24h of flow exposure. **(A)** RNA was harvested from EC exposed to LMMF and the expression of pro-inflammatory genes assessed by qRT-PCR using GAPDH as a housekeeping gene (n=3-4; analysis by unpaired t-test). **(B)** EC were fixed and incubated with anti-VE-cadherin antibody, 488-phalloidin and DRAQ5 nuclear stain (n=4; scale = 50 µm; representative images shown).

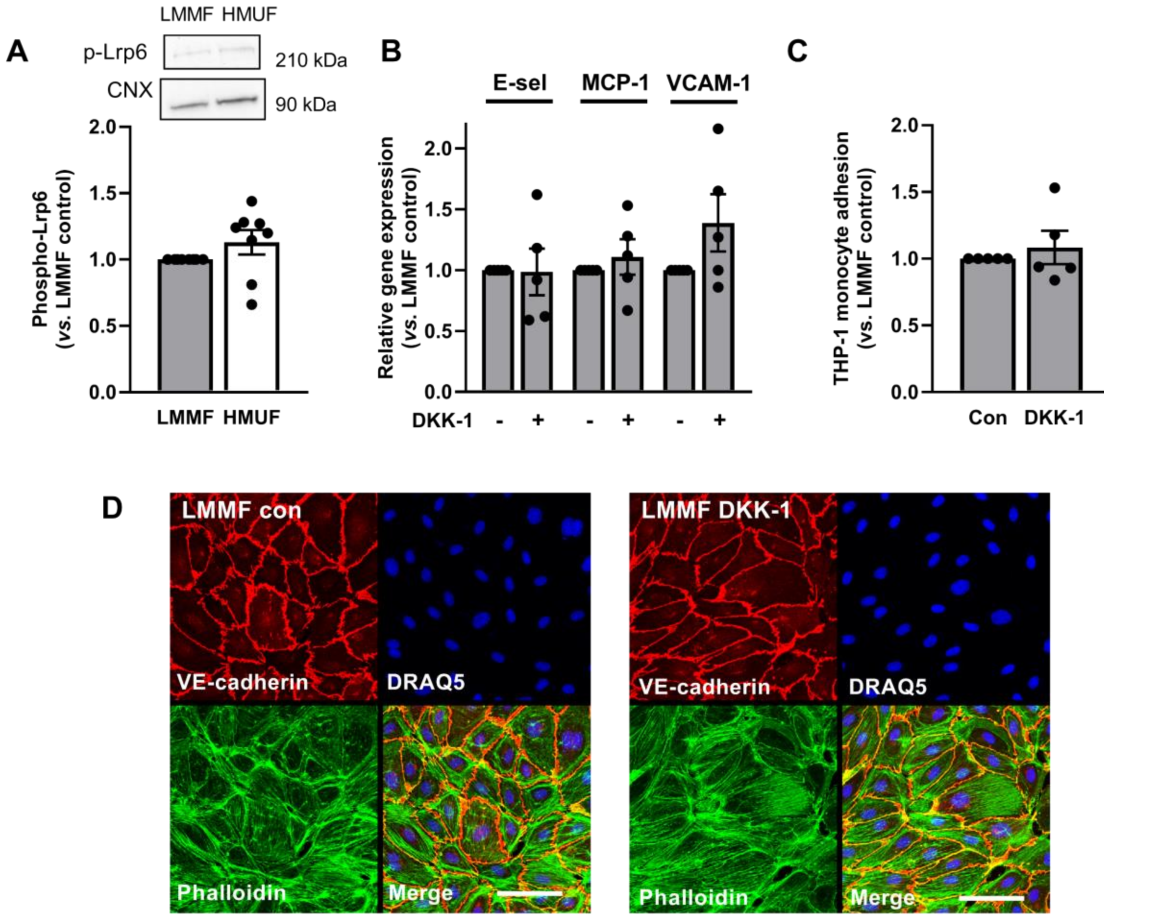


**Supplementary Figure 10. Inhibition of the canonical Wnt pathway using DKK-1 has no effect on the phenotype of EC to LMMF**

**(A)** Lysates were obtained from EC exposed to LMMF and HMUF for 72h and the activation of Lrp6 assessed by western blot using calnexin (CNX) as a loading control (n=8; analysis by paired t-test). **(B-D)** HAEC were exposed to flow for 72h and treated with DKK-1 (250 ng.ml$^{-1}$) f for the last 24h of flow exposure. **(B)** RNA was harvested from EC exposed to LMMF and the expression of pro-inflammatory genes assessed by qRT-PCR using GAPDH as a housekeeping gene (n=4-5; analysis by unpaired t-test). **(C)** The number of adherent calcein-labelled THP-1 monocytes was determined in 4 fields of view per experiment. Results shown relative to untreated controls (n=5; analysis by unpaired t-test). **(D)** EC were fixed and incubated with anti-VE-cadherin antibody, 488-phalloidin and DRAQ5 nuclear stain (n=3; scale = 50 µm; representative images shown).

#### Supplementary information

##### Antibodies

| Target antigen | Species | Source | Catalog # | Application | Dilution |
|---|---|---|---|---|---|
| β-catenin | Mouse | BD Biosciences | 610153 | IF | 1 in 100 |
| β-catenin | Rabbit | Cell Signaling Technology | 9582 | WB | 1 in 3000 |
| β-catenin (active) | Rabbit | Millipore | 05-665 | WB | 1 in 1000 |
| Calnexin | Goat | Santa Cruz | sc-6465 | WB | 1 in 3000 |
| Frizzled-4 | Goat | Santa Cruz | sc-66450 | WB | 1 in 1000 |
| Frizzled-4 | Rabbit | ThermoFisher Scientific | 710731 | IF | 1 in 100 |
| GSK3β | Rabbit | Cell Signaling Technology | 9315 | WB | 1 in 1000 |
| GSK3β(Ser9) | Rabbit | Cell Signaling Technology | 9322 | WB | 1 in 1000 |
| IκBα | Rabbit | Santa Cruz | sc-371 | WB | 1 in 2000 |
| IκBα(Ser32) | Rabbit | Cell Signaling Technology | 2859 | WB | 1 in 1000 |
| Lrp6 | Rabbit | Cell Signaling Technology | 3395 | WB | 1 in 1000 |
| Lrp6(Ser1490) | Rabbit | Cell Signaling Technology | 2568 | WB | 1 in 1000 |
| NF-κB p65 | Rabbit | Cell Signaling Technology | 8242 | IF | 1 in 500 |
| PDHX (E3BP) | Mouse | Santa Cruz | sc-377255 | WB | 1 in 1000 |
| RSPO-3 | Rabbit | Proteintech | 17193-1-AP | WB | 1 in 1000 |
| Ryk | Mouse | R&D Systems | MAB4907 | WB | 1 in 1000 |
| TBP | Rabbit | Cell Signaling Technology | 44059 | WB | 1 in 2000 |
| VCAM-1 | Rabbit | Cell Signaling Technology | 13662 | WB | 1 in 1000 |
| VE-cadherin | Rabbit | Cell Signaling Technology | 2500 | IF | 1 in 500 |
| VE-cadherin | Mouse | BD Biosciences | 555661 | IF | 1 in 500 |
| Vinculin | Mouse | Sigma | V4505 | IF | 1 in 500 |
| Wnt5a | Rabbit | Proteintech | 55184-1-AP | WB | 1 in 1000 |
| ZO-1 | Rabbit | Cell Signaling Technology | 13663 | IF | 1 in 250 |

##### Reagents

| Target antigen | Species | Source | Catalog # | Dilution |
|---|---|---|---|---|
| AlexaFluor-488 Phalloidin | | ThermoFisher Scientific | A12379 | 1 in 40 |
| DRAQ5 | | Biostatus | DR50200 | 5 µM |
| DKK-1 | Human | R&D Systems | 5439-DK-010 | 250 ng/ml |
| SFRP-1 | Human | PeproTech | 120-29 | 200 ng/ml |
| iCRT5 | | Abcam | ab142141 | 50 µM |

##### Cells

| Cell type | Species | Source | Catalog # |
|---|---|---|---|
| Human aortic endothelial cells | Human | Promocell | C-12271 |
| THP-1 monocytes | Human | ATCC | TIB-202 |

#### Primers

| Gene | Forward | Reverse |
|---|---|---|
| *β-CATENIN* | TGCCCTGGCTATGTGAGTTT | TCAAATACCCTGCATAGTACGCT |
| *CLAUDIN-5* | CCTGTGCCACCGCTTTTTG | CAGCACTGTCTCTCTCATCCC |
| *ENOS* | CATCTTCAGCCCCAAACGGA | AGCGGATTGTAGCCTGGAAC |
| *ESAM* | TCACCAACCTTTCGTCTTCCA | CCAGCGTCACATTACATTGGG |
| *E-SEL* | GCTCTGCAGCTCGGACAT | GAAAGTCCAGCTACCAAGGGAAT |
| *FZD4* | GGATGCTCTGTGGCCTTTCT | GGGCATGTGTAGCAGGAAGT |
| *FZD5* | GAGAGACGGTTAGGGCTCG | TTCTCAGCGGAGTGACCC |
| *FZD6* | GGGAACGGTGGGTTAGACG | CTGGGTCAATTACTCGGGGG |
| *FZD7* | GTCGTGTTTCATGATGGTGC | CGCCTCTGTTCGTCTACCTC |
| *GAPDH* | CTATAAATTGAGCCCGCAGCC | ACCAAATCCGTTGACTCCGA |
| *IL-8* | ACCGGAAGGAACCATCTCAC | GGCAAAACTGCACCTTCACA |
| *JAM-A* | ACGGGAAGACACTGGGACATA | GGATGGAGGCACAAGCAC |
| *JAM-B* | AGCAGTAGAGTACCAAGAGGCTA | CTCCGACCCAGTTTCTTCCA |
| *JAM-C* | CAAAATTCAGGGAGACTTGGCG | CCTCACAGCGATAAAGGGCT |
| *KLF2* | TGGGCATTTTTGGGCTACCT | CCCAGTTCCAAGCAACCAGA |
| *MCP-1* | AGGTGACTGGGGCATTGAT | GCCTCCAGCATGAAAGTCTC |
| *PECAM-1* | CACAGATGAGAACCACGCCT | GGCCCCTCAGAAGACAACAT |
| *RNF43* | TTGTTTCACCCCCGTGGATT | TCACTTGGCATTGCTCTTCC |
| *RSPO1* | GCTGGCAAGGACTGGTGTTT | TGGTTGATTGCCTCGACACC |
| *RSPO2* | AACCGATGGAGACGCAGTAA | GTTGACATCGGCTACACCCA |
| *RSPO3* | GTAGGGGAGAAAGCCACCAC | ACCAGGTTCTCTGAGTTAGCA |
| *RSPO4* | CACAATGGAAAGACCTGCGG | GACTCAGAAAGCACCTGGCA |
| *VCAM-1* | GTCTCCAATCTGAGCAGCAA | TGGGAAAAACAGAAAAGAGGTG |
| *VE-CADHERIN* | ATGAGATCGTGGTGGAAGCG | TGTGTACTTGGTCTGGGTGAAG |
| *ZO-1* | GTGGGTAACGCCATCCTCTG | CCATTGCTGTGCTAGTGAGC |
| *ZNRF3* | CTGGGTAGGAGTGGTGAAGC | CTTGCCTAGGACAGTGAGGC |

**Supplementary Table 1.** Details of resources used for experiments
